# A chromosome-level assembly and functional genomic resources for the model annelid *Capitella teleta*

**DOI:** 10.1101/2025.10.31.685816

**Authors:** Billie E. Davies, Paul Gonzalez, Abhinav Sur, Jingcheng Wei, Tom Frankish, Jimena Montagne, Allan M. Carrillo-Baltodano, Kero Guynes, Yan Liang, Rory D. Donnellan, R. Travis Moreland, Sumeeta Singh, Suiyuan Zhang, Tyra G. Wolfsberg, Néva P. Meyer, Elaine C. Seaver, Andreas D. Baxevanis, José M. Martín-Durán

**Affiliations:** School of Biological and Behavioural Sciences, Queen Mary University of London, Mile End Road, E1 4NS, London, United Kingdom; Center for Genomics and Data Science Research, Division of Intramural Research, National Human Genome Research Institute, National Institutes of Health, Bethesda, Maryland 20892, USA; Division of Intramural Research, Eunice Kennedy Shriver National Institute of Child Health and Human Development, National Institutes of Health, Bethesda, Maryland 20892, USA; Blizard Institute, Barts and The London School of Medicine and Dentistry, QMUL, London, E1 2AT, UK; Wellcome Sanger Institute, Wellcome Genome Campus, Hinxton, CB10 1SA, Cambridge, United Kingdom; Biology Department, Clark University, Worcester, Massachusetts, USA; Whitney Laboratory for Marine Bioscience, University of Florida, 9505 Ocean Shore Dr., Saint Augustine, FL 32080, USA

**Author notes:** Correspondence: Elaine C. Seaver, Andreas D. Baxevanis, José M. Martín-Durán.

**Keywords:** *Capitella teleta*, Annelida, Spiralia, genome, genome browser, mitochondrial genome, chromosome, macrosynteny, comparative genomics, single-cell RNA-seq

## Abstract

**Background:** The polychaete *Capitella teleta* is a commonly used annelid for studies in evolutionary developmental biology, comparative genomics, conservation, and ecotoxicology. Over a decade ago, it was the first polychaete to have its genome sequenced and assembled, contributing to pioneering studies that transformed our understanding of animal genomes and their evolution. However, this early resource is now outdated compared to current genome sequencing standards, limiting the use of modern functional genomic tools that could further our understanding of numerous biological processes.

**Results:** We combine long-read and short-read sequencing with Hi-C chromatin conformation capture data to assemble the chromosome-level nuclear and mitochondrial genomes of the laboratory strain of *C. teleta*. This reference assembly more accurately reflects the expected genome size for this polychaete (∼243.6 Mb) and contains a highly complete, evolutionarily conserved gene repertoire. Notably, the nuclear and mitochondrial genomes are heavily rearranged, indicating a decoupling between gene family repertoire and chromosomal evolution. The analyses of multi-omic datasets available for *C. teleta*, including developmental time courses of bulk and single-cell RNA-seq, ATAC-seq, and EM-seq, using the new reference assembly, resulted in a significant quality improvement, allowing us to identify new cell-type-specific gene markers and gain additional insights of biological relevance. Finally, we generated a publicly available genome browser that ensures all these resources are easily findable, accessible, interoperable, and reusable.

**Conclusions:** Our study provides state-of-the-art genomic resources for the polychaete model *C. teleta*, addressing a pressing community need that will open new research opportunities in animal and genome evolution.

## Background

Chromosome-level reference genome assemblies are the current gold standard in genomics and an indispensable resource for most areas of biological research. Although traditionally restricted to a small number of biomedical model systems, such as humans, mice, *Drosophila,* and *C. elegans*, technical and cost-cutting advances have transformed the production of chromosome-level assemblies, enabling ambitious initiatives aimed at generating reference genome assemblies for a sizable fraction of the planet’s biodiversity [1, 2]. Despite these efforts, taxonomic gaps still persist, especially within Spiralia, one of the three major lineages of bilaterally symmetrical animals [3], and among the species that pioneered the field of comparative genomics over a decade ago [4]. Moreover, most genomes resulting from large sequencing initiatives lack dedicated, user-friendly platforms that facilitate the exploration, integration, and reuse of genomic information, thereby limiting the impact of these datasets on the research community.

*Capitella teleta* (Blake, Grassle, & Eckelbarger, 2009; formerly known as *Capitella* sp. I) is a marine polychaete worm (Annelida) that has been widely used as a model in developmental and regenerative biology, comparative genomics, conservation, and ecotoxicology (Figure 1A) [5]. With a clearly segmented body, *C. teleta* exhibits a burrowing lifestyle. Like many annelids, this worm continues to generate new segments throughout its life and can regenerate all somatic and germline tissues along its primary body axis posteriorly [6]. Every segment of *C. teleta*’s body is morphologically complex. It contains a ganglion of the centralised nervous system, body wall musculature, gut, epidermis, bristles and peripheral sensory systems. Additionally, a subset of segments contains gonads and nephridia. Initially cultured as a laboratory strain over 50 years ago due to its ability to thrive in disturbed habitats, *C. teleta* is among the best-studied of the over 18,000 annelid species described to date, providing key insights into its molecular development and regeneration [7].

**Figure 1.**
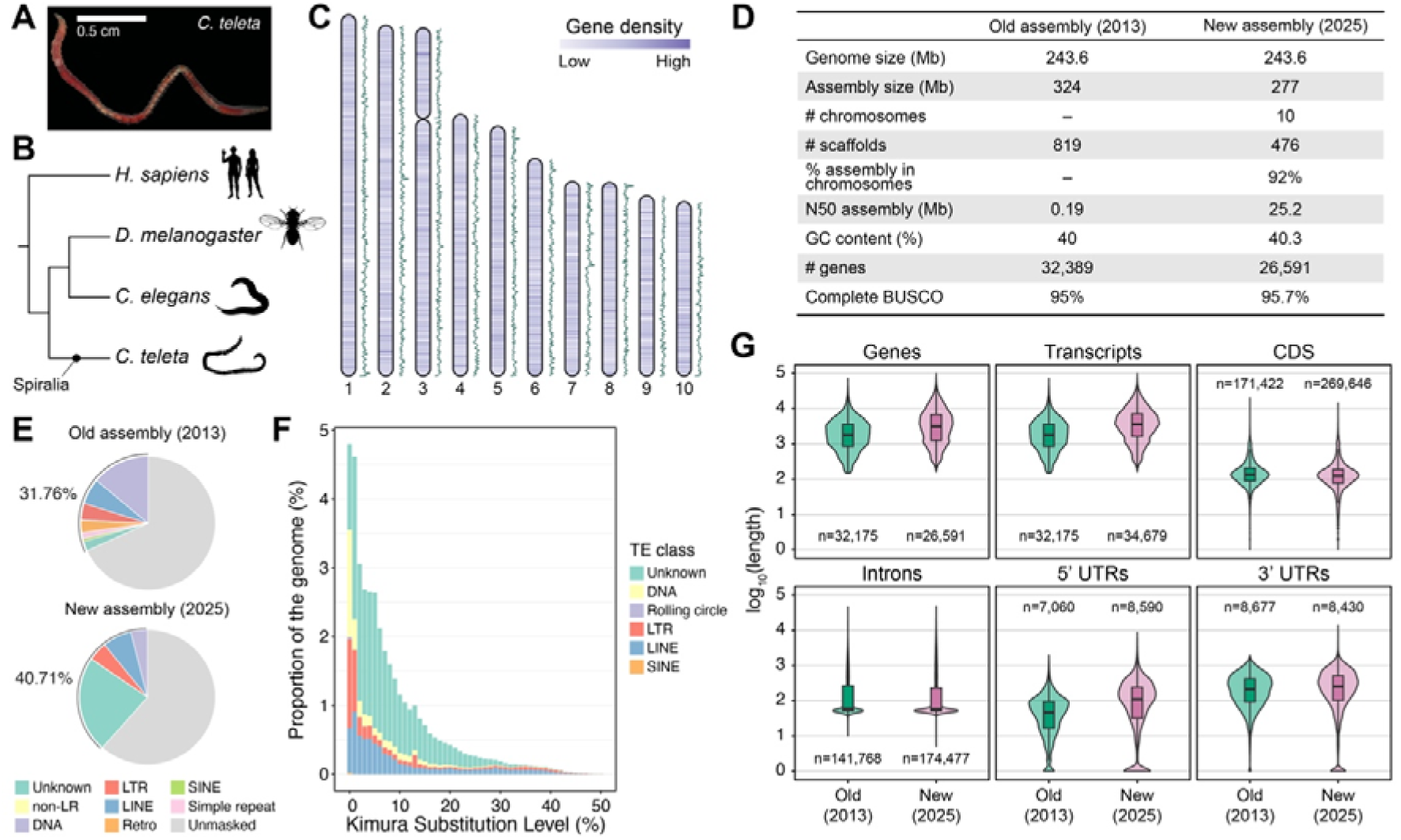
A chromosome-level genome assembly and updated annotation of the annelid *Capitella teleta*. (**A**) An adult *C. teleta* specimen. Scale bar, 0.5 cm. (**B**) Phylogenetic tree showing the evolutionary position of *C. teleta* (a spiralian) relative to common model organisms. (**C**) Ideogram of the 10 pseudochromosomes in the new assembly, with shading indicating gene density and the green line indicating transposable elements (TEs) density. (**D**) Table comparing key metrics of the old (2013; [4]) and new (2025; this study) genome assemblies, highlighting significant improvements in contiguity (N50 increased from 0.19 Mb to 25.2 Mb) and contiguity (92% of the assembly is anchored to pseudochromosomes). (**E**) Pie charts comparing the annotated repeat content of the old and new assemblies. The new assembly identifies a larger proportion of the genome as repetitive elements (40.71% vs 31.76%). (**F**) Transposable element (TE) landscape of the new assembly. The abundance of different TE classes is plotted against their Kimura substitution level, indicating bursts of TE activity (peak at low divergence) that are progressively removed. (**G**) Violin plots comparing the length distributions of annotated genes and gene features between the old and new assemblies. The new assembly resolves significantly longer genes, transcripts, and introns, while the lengths of coding sequences (CDS) and untranslated regions (UTRs) remain comparable. Sample sizes (n) for each feature are indicated on the plots.

*Capitella teleta* is a member of Spiralia, one of the three superclades of bilaterian animals (Figure 1B) [8]. Its highly stereotyped cleavage program has enabled a cell lineage-based approach to determine the embryonic source of every cell in the adult animal, resulting in the generation of a comprehensive fate map [9, 10]. Following embryogenesis, the life cycle of *C. teleta* includes the formation of a nonfeeding larva before metamorphosing into a juvenile burrowing worm. The year-round availability of embryos in laboratory cultures, their synchronised development, and an established developmental staging system have all greatly facilitated the establishment of *C. teleta* as a tractable experimental system [11]. Published functional genomic methods, such as CRISPR/Cas9 mutagenesis, morpholino knockdown, exogenous RNA expression, and transcriptomic profiling, as well as experimental techniques such as blastomere and juvenile microinjections, microsurgery, and single blastomere deletions, have been successfully applied [6, 12–18]. As a result, the genome of *C. teleta* was the first polychaete genome and one of the first spiralian genomes to be sequenced [4]. While existing genomic data have greatly facilitated comparative genomic studies and provided an invaluable resource for polarising evolutionary genomic events in the Bilateria, the field would greatly benefit from highly contiguous whole-genome sequence data. Such a resource would enable researchers to thoroughly address key biological processes like animal whole-body regeneration, chromosome structure and evolution, cis-regulation of gene expression, and epigenomics.

Here, we combine long- and short-read sequencing with chromatin conformation capture scaffolding to produce a *de novo* chromosome-level genome assembly for the annelid *C. teleta*. Our assembly accurately resolves the 10 chromosome pseudomolecules inferred from karyotypic analyses [19], resulting in a highly complete genome assembly that aligns more closely with the estimated genome size (243.6 Mb) than that reported in the original sequencing effort [4]. Using extensive transcriptomic evidence [17, 18], we annotated 26,591 protein-coding genes and confirmed that *C. teleta* exhibits a slow-evolving gene repertoire, having retained a large proportion of ancestral metazoan gene families. We generated the first assembly of the mitochondrial genome for the lab strain (derived from the holotype) of *C. teleta*, which, like the nuclear genome, has experienced profound macrosyntenic rearrangements. The use of our new chromosome-level assembly also improves the analyses of available functional genomic datasets, including developmental time course data of transcriptomic (bulk and single-cell) expression, determination of open chromatin regions, and DNA methylation profiling [16–18, 20]. Indeed, the new genome annotation captures more cell barcodes and resolves more clusters in single-cell transcriptomic studies, identifying previously unknown cell-type-specific markers and more granular ectodermal subpopulations. Together, our study provides a reference and cutting-edge genomic resource for the polychaete annelid *C. teleta* while also establishing a publicly accessible genome browser that integrates all available genomic datasets to support and expand the diverse research community studying this invertebrate model system and enable more comprehensive comparative genomic studies.

## Results and Discussion

### A chromosome-level genome assembly of C. teleta

To generate a chromosome-level genome assembly of *C. teleta*, we sequenced 258.21 M reads (∼103 Gb total bases; 372x coverage) Oxford Nanopore long-reads in six different libraries and 194.67 M paired-end reads (∼58.4 Gb total bases; ∼210x coverage) of short Illumina reads (Supplementary Table 1). After assembly, scaffolding, dehaploidisation, and decontamination, we used ∼60 M paired-end reads of HiC data to combine 92% of the assembly length into 10 pseudochromosome scaffolds (Figure 1C; Supplementary Figure 1A, B). Although chromosome 3 is not fully resolved and remains divided into two large scaffolds, the inferred chromosome count matches the karyotype of *C. teleta* [19]. The resulting assembly size is 277.13 Mb — 47 Mb shorter than the original assembly — and is closer to the estimated genome size of *C. teleta* based on *k*-mer distribution analyses (243.6 Mb) (Figure 1D; Supplementary Figure 1C). As expected for an inbred line that has been in closed culture for over 50 years, the levels of heterozygosity are low (0.39–0.41%) compared to wild populations in other annelids (e.g., 2.86–2.9% heterozygosity in *Owenia fusiformis*) [18] (Supplementary Figure 1C). This chromosome-level assembly exhibited a modest improvement in genome completeness compared to the previous one, as inferred through BUSCO searches using the metazoan database (metazoan_odb10; 95.7% vs 95% completeness, respectively; Figure 1D). Indeed, the original and new genome assemblies align well (Supplementary Figure 2A). Collectively, the *de novo* resequencing of *C. teleta*’s genome with long reads and chromatin conformation-based approaches has improved overall contiguity and removed haplotypic redundancy, resulting in an assembly that better matches the inferred genome length and karyotype in this annelid.

The new chromosome-level assembly also improved the annotation of repeats and protein-coding genes. Repeat elements now represent 40.71% of the chromosome-level assembly, an increase from 31.76% in the original version [4] (Figure 1E; Supplementary Table 2). However, the age distribution of transposable elements (TEs) remains equivalent across both assemblies (Figure 1F), supporting the proposition that *C. teleta* has an active repertoire of TEs whose expansions are progressively purged from the genome, despite not exhibiting obvious signs of TE-associated DNA methylation [16]. We generated *de novo* functional annotation using the extensive transcriptomic resources recently generated for *C. teleta* [17, 18], predicting 26,591 protein-coding genes (Figure 1D), 5,798 fewer than in the original assembly [4]. This reduction is consistent with a smaller assembly length and the removal of redundant haplotypes. Yet, both annotations are broadly similar, as indicated by the high proportion of complete BUSCO genes in both versions, as well as the 17,797 genes (66.93%) in the new annotation that have a one-to-one best reciprocal BLAST hit with a gene model in the original assembly (Supplementary Figure 2B). As expected, the increase in assembly contiguity has resulted in longer gene and transcript lengths, primarily driven by improved UTR annotations (Figure 1G). Additionally, a larger proportion of transcripts now have BLASTX hits against the UniProt database, as well as Gene Ontology and KEGG annotations (Supplementary Figures 2C, D), indicating that more genes in the new assembly can be confidently assigned a biological function. Thus, our new chromosome-level assembly for *C. teleta* provides a substantially enhanced, high-quality reference genome that will be invaluable for future functional and comparative studies.

### A slow-evolving gene family repertoire in a fast-evolving karyotype

Given the notable reduction in gene number between the original and chromosome-scale assemblies (Figure 1D), we performed gene-family evolution analyses to identify gene gains and losses in *C. teleta*. For this purpose, we collated a dataset of 31 chromosome-level assemblies, covering all major bilaterian groups, with an expanded sampling within Annelida (Supplementary Tables 3 and 4). While the last common ancestor of Spiralia gained gene families, gene losses outweighed gains in the evolution of Annelida (Figure 2A; Supplementary Figure 3). Notably, predominant patterns of gene loss are also observed at the nodes corresponding to the evolution of major internal groups within Annelida, including Sedentaria, Errantia, Siboglinidae, and Clitellata (Figure 2A; Supplementary Figure 3). Despite being a member of Sedentaria, *C. teleta* has retained a comparable number of ancestral Eumetazoan and Bilaterian orthogroups compared to other slow-evolving annelids, such as *O. fusiformis*, and even more than what has been observed in humans (Figure 2B) [18]. Additionally, the improved gene annotation is reflected in ortholog assignment, with the new chromosome-level assembly exhibiting a higher proportion of orthologs (including one-to-one orthologs) with the human genome (Figure 2C). Therefore, the new set of annotations confirms that this polychaete harbours a highly conserved, relatively slow-evolving gene repertoire.

**Figure 2.**
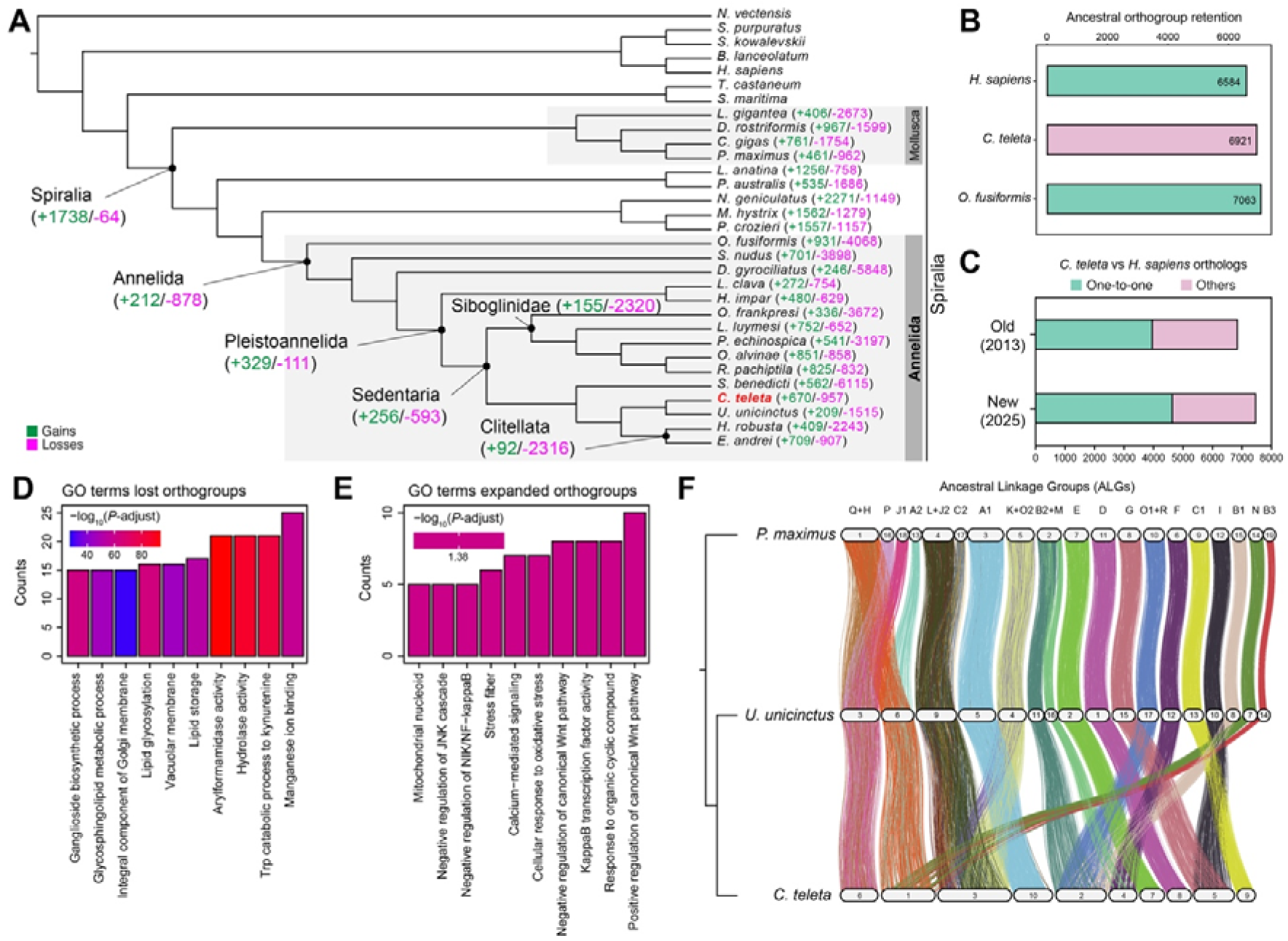
The evolution of the gene family repertoire in *C. teleta*. (**A**) Cladogram of representative bilaterians with increased sampling within Annelida. Numbers at key nodes indicate the inferred number of gene family gains (green) and losses (magenta). (**B**) Comparison of ancestral orthogroup retention between the annelids *C. teleta* and *O. fusiformis* and the human (*Homo sapiens*). (**C**) The new annotation identifies substantially more one-to-one orthologs between *C. teleta* and *H. sapiens* than the original assembly. (**D**, **E**) Gene Ontology (GO) term enrichment for orthogroups lost (**D**) and expanded (**E**) in *C. teleta*. Lost genes are enriched for metabolic processes, while expanded genes are associated with stress and immune response, as well as developmental signalling pathways. (**F**) Conserved synteny between the annelids *Urechis unicinctus* and *C. teleta*, and the mollusc *Pecten maximus* (as a close representative of the ancestral state). Unlike *U. unicinctus* and other polychaetes, *C. teleta* has experienced multiple fusions and fissions of the ancestral linkage groups. Lines connect orthologous genes, and coloured blocks represent ancestral linkage groups.

To explore the implications of the gains and losses of gene families in *C. teleta*, we performed Gene Ontology (GO) term enrichment on significantly gained and lost orthogroups in this annelid (Supplementary Table 5). Most of the lost gene families are involved in metabolism (Figure 2D), ranging from manganese ion binding and catabolism to kynurenine, a tryptophan metabolite involved in diverse biological functions [21], which is the most highly represented. Interestingly, the expanded gene families include those involved in developmental processes, primarily the Wnt, EGFR, and JNK signalling cascades, as well as neurodevelopment (e.g., the secreted extracellular matrix protein SLIT) and the immune response (NF-kB pathway) (Figure 2E; Supplementary Table 6). Previous analyses identified G-protein-coupled receptors (GPCRs) to be highly expanded in *C. teleta* [4, 22]. Consistent with these findings, we also identified five expanded orthogroups containing at least one gene annotated as a GPCR, totalling 214 genes (Supplementary Table 6). The expansion pattern varied, wherein two of these orthogroups were uniquely expanded in *C. teleta* and two others were shared with *Urechis unicinctus* (the sister lineage to *C. teleta*) [23] (Supplementary Table 5). Therefore, the findings confirm the expansion of GPCRs in this annelid lineage. Considering some of these orthogroups may represent ancestral expansions, further investigation is warranted to assess how this expanded GPCR repertoire may have contributed to the diversification of annelid nervous systems and chemosensation.

Some of the most extensive chromosomal rearrangements in animals occur in Annelida [24–27]. The original analysis of ancestral linkage group conservation (ALG) in *C. teleta* revealed signatures of macrosynteny retention [4]. However, *C. teleta* has 10 chromosomes, and the estimated ancestral chromosome number for Annelida is 19 [24], indicating that extensive fusion events must have occurred in this annelid. To resolve the differences in ancestral gene linkage in *C. teleta*, we compared the chromosomal location of one-to-one orthologs between *C. teleta* and *U. unicinctus*, as well as the bivalve mollusc *Pecten maximus* and the nemertean worm *Lineus longissimus* (two annelid outgroups with well-conserved ALGs; Figure 2F; Supplementary Figure 4A–C). As hypothesised, numerous fusion and fission events have occurred in *C. teleta* (Figure 2F; Supplementary Figure 4D). Compared with the ancestral condition for Pleistoannelida (Errantia + Sedentaria) [24], the *C. teleta* genome has undergone 10 ALG fusions, as well as the fission of one chromosomal arrangement (H+Q) present in the last common annelid ancestor (Figure 2F; Supplementary Figure 4D). Notably, the fission of H+Q and its subsequent fusion with A2 and P+J1, respectively, is also present in *U. unicinctus* [24] and may represent a shared synapomorphy for the clade rather than *C. teleta*-specific divergences. Therefore, our findings indicate that ALG evolution in *C. teleta* mirrors that of *O. fusiformis*, a slow-evolving annelid with a well-conserved gene repertoire and numerous chromosomal rearrangements [18, 24]. This suggests that chromosomal, gene family, and molecular evolution rates are uncoupled in several lineages within Annelida, thereby raising caution about the significance and usefulness of ALG conservation in phylogenetic studies [28–30].

### The mitochondrial genome of C. teleta

Despite *C. teleta*’s relevance in evo-devo and ecotoxicology studies, a mitochondrial genome assembly for this species–– particularly for the laboratory culture population broadly used in experimental analyses–– has heretofore been unavailable. To fill this gap, we used Illumina short-read data to generate a complete, circularised mitochondrial genome assembly. This genome has two replication origins, a generally positive GC skew, and a complete gene set comprised of 13 coding genes, two rRNAs, and 22 tRNAs (including two tRNAs for methionine, M0 and M1) (Figure 3A). This gene content is similar to that observed in other annelid mitochondrial genomes [31, 32]. Although most genes (8/13; 61.5%) use the standard ATG start codon, alternative start codons such as ATT (3/13; 23.1%), ATA (1/13; 7.7%; in *nad6*), and the relatively rare TTA (1/13; 7.7%; in *cox1*) also occur. Notably, TTA in *cox1* is also found in some insects [33]. The species description of *C. teleta* reported the holotype’s *cox1* sequence [5], which appears to be profoundly divergent but still closely related to the one reported here (Supplementary Figures 5 and 6; Supplementary File 1). Indeed, the phylogenetic analysis of the publicly available *cox1* sequences for the genus *Capitella* revealed a complex phylogenetic pattern. *Capitella capitata* and *C. neociculata* likely represent species complexes [34], while *C. nonator* from Brazil and *C. teleta* (cosmopolitan, including the holotype and the *cox1* sequence reported here) are well-defined monophyletic clades (Supplementary Figures 6 and 7). These results underscore the need for a comprehensive reassessment of the genus *Capitella* that incorporates additional mitochondrial and nuclear genes alongside morphological descriptions [35–37].

**Figure 3.**
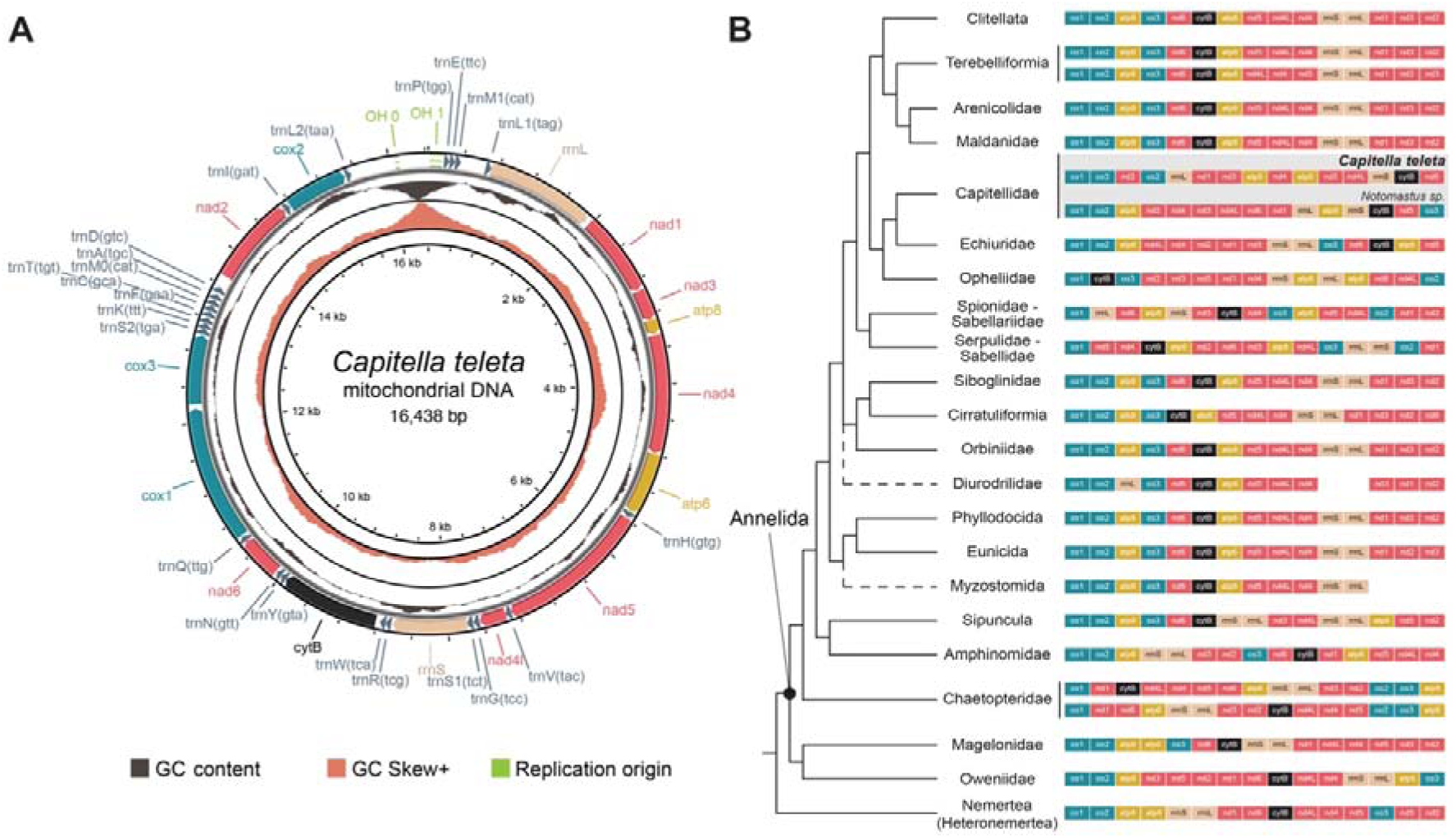
The mitochondrial genome of *C. teleta* and comparative gene order in Annelida. (**A**) Gene map of the 16,438 bp mitochondrial genome of *C. teleta* (lab strain). Genes are shown on the outer ring. Inner rings depict GC content (black) and GC skew (orange). Replication origins are shown in light green. *Capitella teleta*’s mitochondrial genome includes 12 protein-coding genes, two ribosomal RNAs and 23 tRNAs. (**B**) Comparison of mitochondrial gene order across the annelid phylogeny, depicting the mitochondrial gene order of Heteronemertea as an outgroup. The arrangement in Capitellidae (grey box) and in particular *C. teleta*, deviates significantly from the ancestral condition of the last common ancestor to Errantia and Sedentaria.

The mitochondrial genome has also experienced extensive reorganisation during annelid evolution [31, 32] (Figure 3B). While early branching lineages such as Owenidae, Magelonidae, and Chaetopteridae display plastic syntenic arrangements in their mitochondrial genomes when compared to the outgroup Nemertea, it appears that a stable mitochondrial gene order originated in the last common ancestor of Errantia and Sedentaria; it is also possible that this gene order originated even earlier in the last common ancestor of Errantia, Sedentaria, Sipuncula, and Amphinomidae (Figure 3B) [31, 32]. Minor secondary rearrangements have occurred in some lineages of Sedentaria, such as Cirratuliformia. However, extensive changes to the ancestral condition are observed in Spionidae/Sabellariidae, Serpulidae/Sabellidae, and the clade comprising Opheliidae, Echiuridae, and Capitellidae, in which even *C. teleta* and *Notomastus* sp. exhibit contrasting gene orders (Figure 3B) [38]. The gene order of the *C. teleta* lab strain is similar to that recently reported for a *C. teleta* specimen from China [38], which we experimentally validated using diagnostic PCR (Supplementary Figure 8). Collectively, *C. teleta* exhibits highly rearranged nuclear and mitochondrial genomes, unlike other annelid lineages that display dramatically shattered nuclear genomes but stable mitochondrial gene orders (e.g., Clitellata) or relatively conserved ancestral linkage groups but rearranged mitochondrial genomes (e.g., *U. unicinctus*).

### Functional genomic resources for C. teleta

In recent years, methodologies for profiling and interrogating the functional elements controlling the unfolding of genomic information have become available for lesser-studied animals. In *C. teleta*, these methods have been used to perform bulk and single-cell RNAseq profiling throughout its life cycle, specifically during early embryonic cleavage [17, 18].

Additionally, the Assay for Transposable-Accessible Chromatin with high-throughput sequencing (ATAC-seq) has been used to perform time-course experiments aimed at identifying open chromatin regions, while Enzymatic Methyl-seq (EM-seq) approaches have been used to profile temporal changes in 5-methylcysteine (5mC) DNA methylation across the *Capitella* genome (Figure 4A) [16, 18]. Ultimately, the application of these and other genome-wide analyses methods strongly benefit from the availability of highly contiguous genome assemblies and improved annotations. Therefore, we remapped and reanalysed these RNA-seq, ATAC-seq and EM-seq datasets using our new chromosome-level genome assembly and annotations for *C. teleta*. The time course of the bulk RNA-seq profiling experiments spans 15 developmental time points from the oocyte to the adult stage, including each round of cell division during spiral cleavage and most larval stages (Figure 4A). These are highly correlated samples despite having been generated in three different studies, and they cluster in three large groups by principal component analyses: one grouping consists of oocyte and cleavage stages, another group consists of gastrula and early/mid larval stages (i.e., organogenesis), and a final group contains late larval and adult samples (Figure 4B).

**Figure 4.**
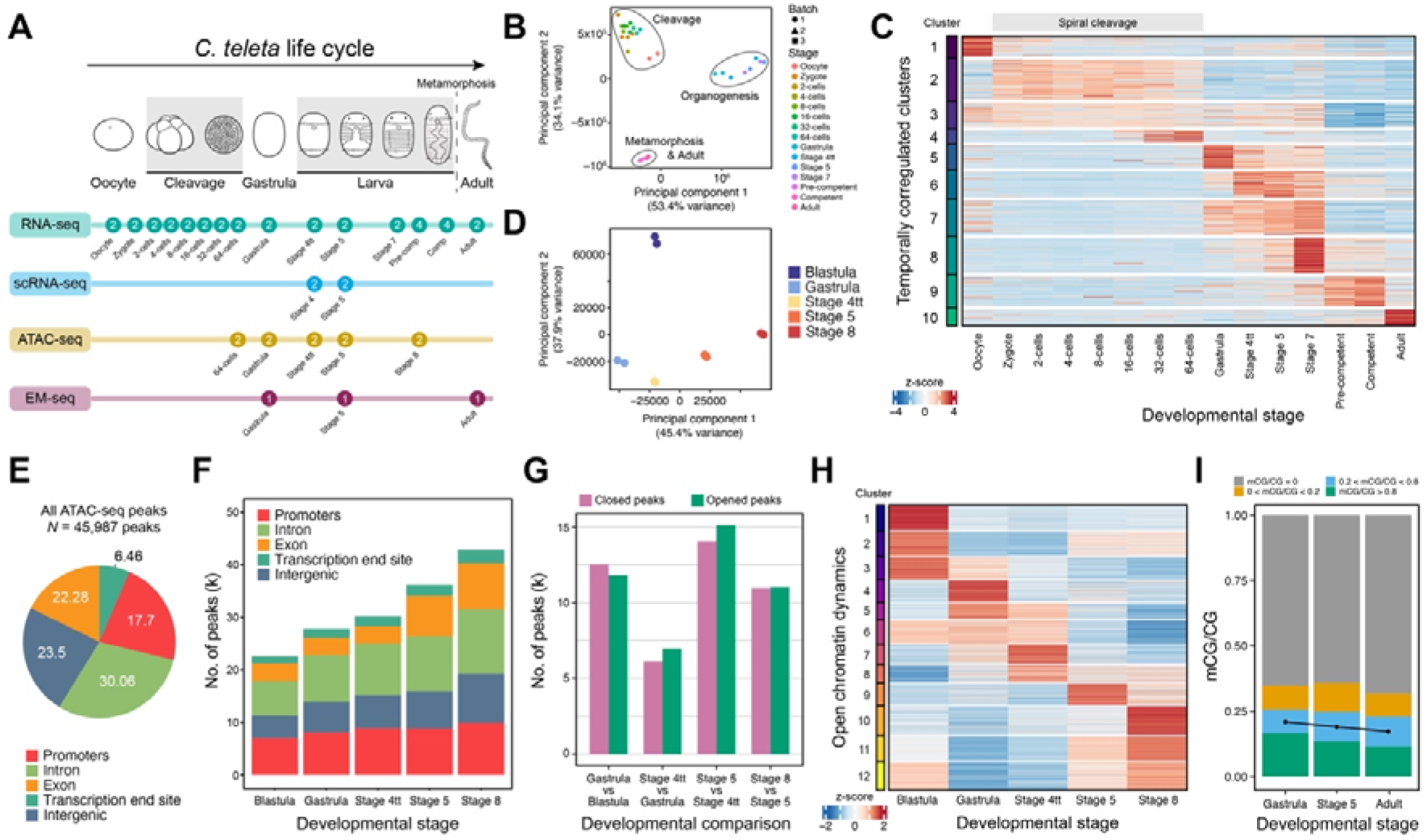
Multi-omic profiling reveals the regulatory dynamics of *C. teleta* development. (**A**) Schematic of the *C. teleta* life cycle, with timelines indicating the stages sampled for RNA-seq, single-cell RNA-seq (scRNA-seq), ATAC-seq, and Enzymatic Methyl-seq (EM-seq). (**B**) A Principal Component Analysis (PCA) of the bulk RNA-seq time course reveals three large transcriptomic groups as the life cycle progresses. (**C**) A heatmap of soft *k*-means clustered bulk RNA-seq data reveals clusters of genes with coordinated expression patterns, indicating distinct temporal waves of gene activation and repression throughout the life cycle. (**D**) PCA of bulk ATAC-seq samples confirms that each developmental stage has a unique chromatin accessibility profile. (**E**) Consensus ATAC-seq peaks distribute more or less equally across intergenic, intronic, exonic, and promoter regions. (**F**) The total number of accessible peaks increases as development progresses, while the genomic annotation of these peaks remains constant. (**G**) Bar plots showing dynamic opening and closing of chromatin, with the largest reshaping of the regulatory landscape happening between stages 4tt and 5, at the beginning of organogenesis. (**H**) A heatmap of soft *k*-means clustered peaks based on their accessibility reveals groups of temporally coregulated regulatory elements. (**I**) Bar plots indicating the proportions of DNA methylation levels at three time points of the life cycle, supporting a global demethylation in adulthood.

This high-resolution dataset profiles the expression of 34,333 transcripts throughout the life cycle, grouped into 10 clusters of temporally coregulated genes by soft *k*-means clustering (Figure 4C). Clusters 1 to 4 are primarily restricted to cleavage stages (from oocyte to the 64-cell stage), while clusters 5 to 8 include genes upregulated from gastrula to the mid-late larval stage (stage 7) (Figure 4C). Finally, clusters 9 and 10 comprise genes that are more highly expressed in the competent and adult stages; these genes may thus be involved in metamorphosis and the establishment of the definitive adult characters of the annelid *C. teleta* (Figure 4C). Accordingly, clusters 1 to 4 (cleavage) are enriched in Gene Ontology (GO) and Kyoto Encyclopaedia of Genes and Genomes (KEGG) terms related to metabolism and cell cycle regulation, while clusters 5 to 8 (gastrulation and larval development) are enriched in gene expression (transcription and translation), cilia development and neurogenesis GO terms, all of which are consistent with the main developmental processes occurring at these stages of the *C. teleta* life cycle (Supplementary Figures 9 and 10). Finally, GO terms in clusters 9 and 10 (competent larvae and adults) are enriched for external stimuli processing and immunity (Supplementary Figure 9). Thus, this high-resolution transcriptomic time course serves as a comprehensive resource for exploring the temporal orchestration of essential developmental genetic programmes in this emerging spiralian model.

Chromatin accessibility, especially in promoter regions, and 5mC levels correlate with gene expression in *C. teleta* [16, 18], making them two essential sources of genome regulatory data for investigating gene expression control. As expected, a chromosome-level assembly improved read mapping for both ATAC-seq and EM-seq datasets. The increase in contiguity and the removal of haplotypic blocks also increased the number of statistically consistent open chromatin regions across the five highly correlated ATAC-seq data points (44,368 vs 45,897 IDR peaks in the old and new genome assembly, respectively) (Figure 4E) [18].

Consistent with previous analyses [18], the 45,897 ATAC-seq peaks are distributed relatively evenly across the genome (Figure 4E), with a gradual increase in peak numbers as development progresses (Figure 4F). Pairwise comparisons of sequential time points revealed thousands of differentially accessible open chromatin regions (p-value < 0.05), with the transition from stage 4tt to stage 5 exhibiting the most considerable change in the open chromatin landscape (Figure 4G). This is consistent with the onset of critical developmental processes at this early larval stage, including the development of the nervous and muscle systems, segments, and eyes [39, 40]. Notably, soft *k*-means clustering of accessibility dynamics defined 12 clusters of temporally co-regulated open chromatin regulatory elements, five of which (clusters 1, 4, 7, 9 and 10) were mainly stage-specific (Figure 4H). Finally, the increased mappability of EM-seq data also improved the detection of 5mC levels across the life cycle, supporting the gradual decrease in global DNA methylation levels as development and metamorphosis progress in this annelid (Figure 4I) [16]. Taken together, these analyses reinforce the benefits of a chromosome-level assembly, which can serve as a reference resource for exploring how epigenomic data influence gene expression and development in *C. teleta*.

*The new assembly improves the analysis of single-cell RNA-seq data in* C. teleta Single-cell RNA sequencing (scRNA-seq) is increasingly being applied to spiralians, including *C. teleta*, where it has been used to explore cell-type composition in early-stage larvae at 24- and 48-hour post-gastrulation (stages 4 and 5) [20]. In most cases, scRNA-seq workflows rely on 3’ end-sequencing to identify and count transcripts in each cell as a measure of gene expression. Thus, we reasoned that the improvements in assembly contiguity and gene annotation (Figure 1D, G) with increased 3’ UTR coverage would improve scRNA-seq data analyses. To test this, we combined and re-analysed previously published scRNA-seq datasets, quantified reads mapped to the old and new annotations using the CellRanger pipeline, and then filtered, clustered, and performed differential expression analysis using Seurat [41, 42]. For cell clustering, we utilised the first 14 components, which contribute to a comparable amount of cumulative variation in both cases (Supplementary Figure 11A, B). Initial CellRanger mapping revealed an increase in the number of cells captured, but a lower read depth per cell and a lower median number of UMIs/cell (Supplementary Figure 11C) when using the new annotation compared to the old assembly. However, most other standard metrics were comparable between the two annotations. Upon combining both stages, we identified nine clusters from 12,745 cells and 11 clusters from 17,115 cells when mapping to the old and new annotations, respectively (Figure 5A; Supplementary Figures 12–22). We also observed an increase in the total number of genes detected using the chromosome-level assembly and annotation (Figure 5A). This suggests that the improved 3’ UTR coverage and assembly completeness increased cell and gene calls used for clustering.

**Figure 5.**
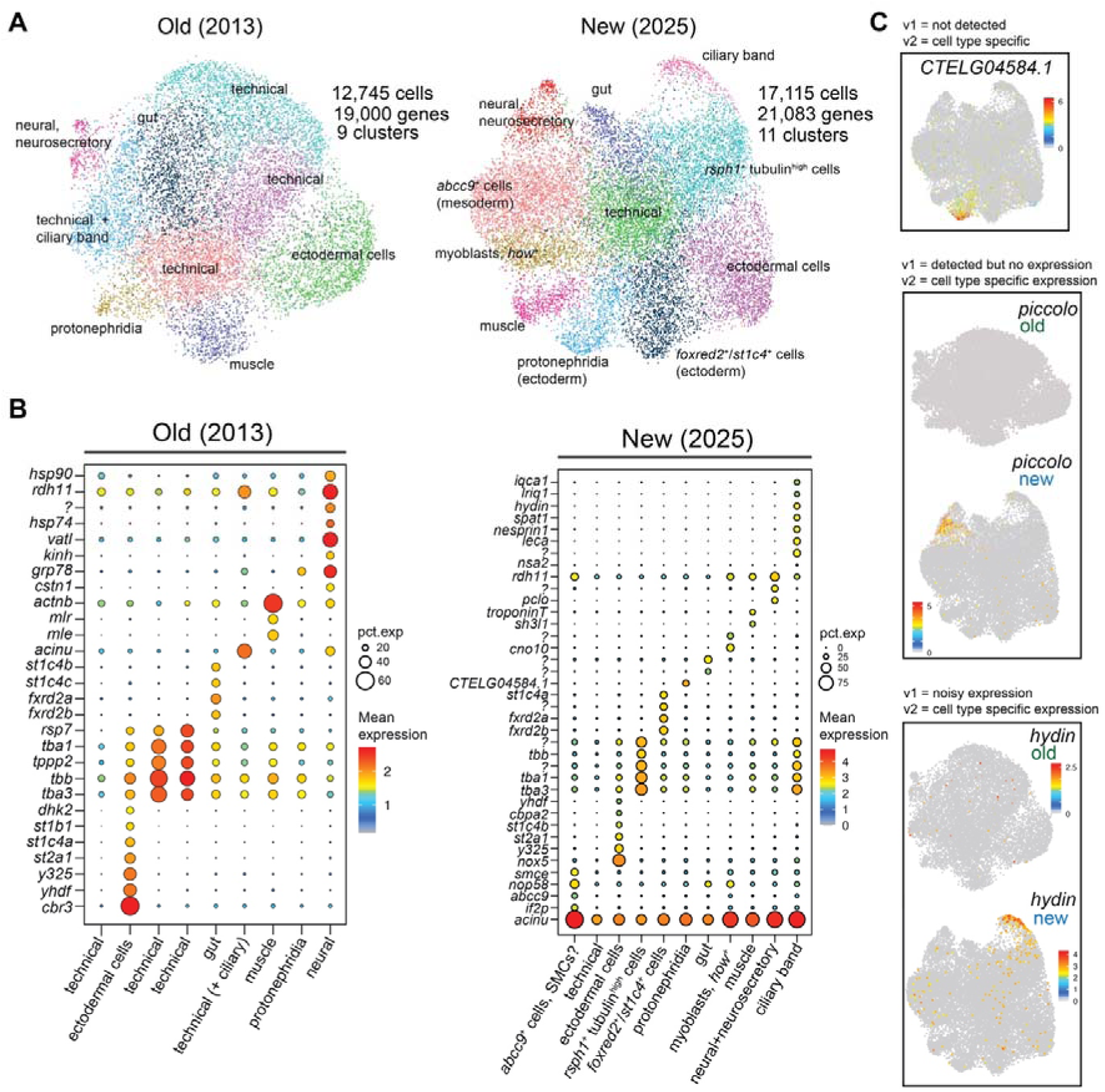
A chromosome-level assembly enhances the analysis and resolution of single-cell transcriptomics. (**A**) UMAP visualisations of single-cell RNA-seq data from stage 4 and 5 larvae, analysed with the old [4] and new (this study) genome annotations. The new annotation identifies more cells (17,115 vs. 12,745) and resolves more distinct cell clusters (11 vs. 9), including the separation of ectodermal and mesodermal subpopulations. (**B**) Dot plots comparing the expression of key marker genes across clusters. The new annotation offers higher-resolution expression patterns, enabling a more precise definition of cell types. Dot size corresponds to the percentage of expressing cells, and the colour indicates the mean expression level. (**C**) UMAP plots illustrating how the new annotation (v2) enables the detection and localisation of genes that showed noisy or no expression in the old annotation (v1). For example, *CTELG04584* is a novel marker of protonephridia, *piccolo* expression is now clearly localised to neural clusters, and *hydin* expression is specific to the ciliary band.

The increased number of cell clusters after using the new annotation suggests that some previously identified clusters may have resolved granularly due to the greater number of genes and cells. This is further supported by the lower number of technical clusters (i.e., the lack of cluster-specific genes) using the chromosome-level assembly annotation (Figure 5A, B). To further compare the outputs from the two annotations, we identified three cell clusters (ectoderm, neural, and ciliary bands) that could be unambiguously matched between the old and new analyses based on cluster-specific gene expression signatures in each dataset (Figure 5A, B). Among these three clusters, we were able to detect genes that could be categorised into three groups: (i) not detected in the old dataset but showing cell type-specific expression in the new one (e.g., *CTELG04584.1*); (ii) detected in both datasets, but expressed in a cell type-specific manner using the new assembly while not expressed at discernible levels in the original analysis (e.g., expression of *piccolo*, a presynaptic protein [43], in the neural cluster); and (iii) noisy expression in the old annotation but cell type-specific expression in the new analysis (e.g., expression of *hydin*, a component of motile cilia [44], in the ciliary bands cluster) (Figure 5C). In this third category, the scRNA-seq analysis using the new annotations detected 318 cells in a ciliary band cluster. In contrast, the transcriptional profile of ciliary band cells was found to be noisy and categorised as a “technical” cluster in the original analyses (Figure 5A, B) [20]. Within the ciliary band cluster, the new analysis detected previously missed cell type markers, such as *hydin* and *synaptic nuclear envelope 1* (*syne1*), which encodes multiple isoforms of *nesprin1* (nuclear envelope spectrin 1) that localise in the ciliary rootlets [45] (Figure 5B, C). Indeed, *hydin* is expressed in the ciliary bands of stages 5 and 7 larvae (Figure 6A–C), thereby validating these new findings and our cluster annotation.

**Figure 6.**
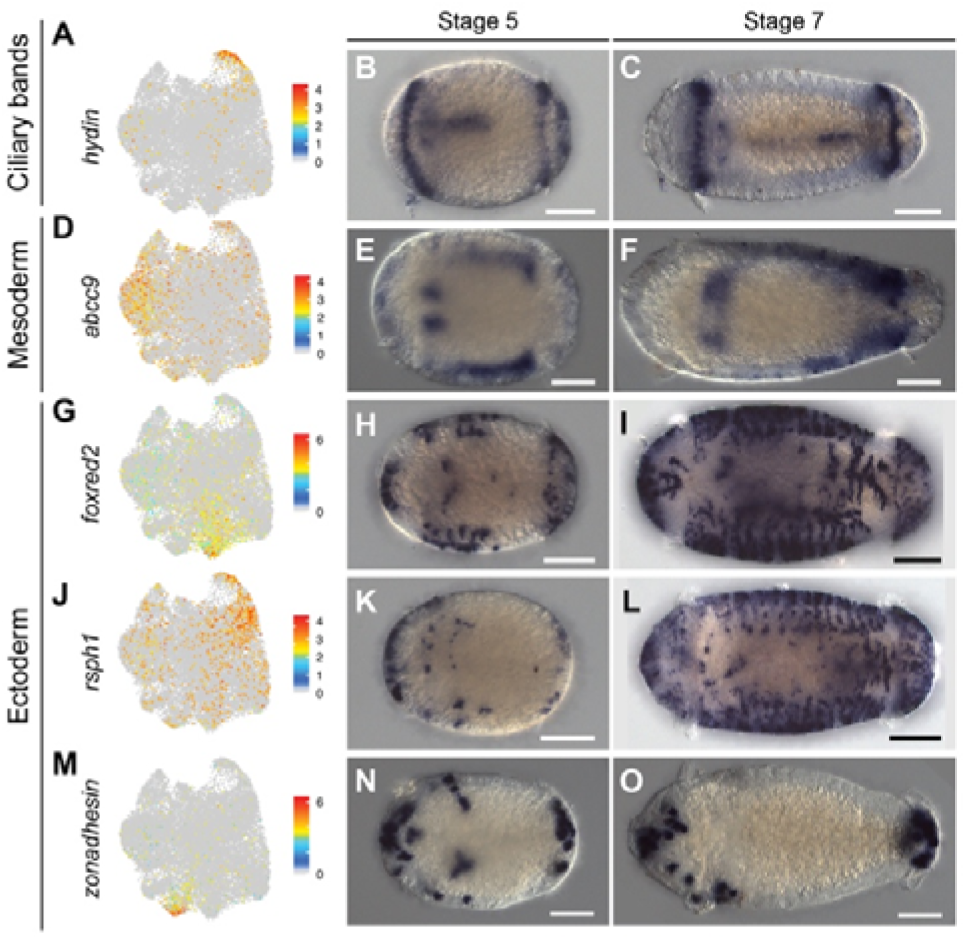
The expression of novel cell-type markers in *C. teleta*. (**A**, **D**, **G**, **J**, **M**) Two-dimensional representation of the scRNA-seq data for stages 5 and 7 larvae, indicating the expression of the ciliary band marker *hydin* (**A**), the mesodermal marker *abcc9* (**D**), and the ectodermal markers *foxred2*, *rsph1*, and *zonadhesin* (**G**, **J**, **M**, respectively). (**B**, **C**, **E**, **F**, **H**, **I**, **K**, **L**, **N**, **O**) Whole mount *in situ* hybridisation of the cell-type-specific markers *hydin*, *abcc9*, *foxred2*, *rsph1* and *zonadhesin* in stage 5 and stage 7 larvae. *Hydin* is expressed in the ciliary bands of the early (**B**) and late (**C**) larvae. The marker *abcc9* is expressed in the lateral mesodermal bands of the early (**E**) and late (**F**) larvae, as well as the mesoderm around the foregut/oesophagus. The markers *foxred2* and *rsph1* are expressed broadly in the ectoderm of both early (**H**, **K**) and late (**I**, **L**) larvae, albeit with different patterns, supporting that these two markers represent different populations, as suggested by the scRNA-seq data (**G**, **J**). The gene *zonadhesin* is expressed in large ectodermal cells at the anterior and posterior ends, as well as ventrally near the mouth, in both early (**N**) and late (**O**) larvae. Larval images are shown in ventral view with anterior to the left. Scale bars are 50 µm.

Among the granular clusters captured in the new dataset, there was unexpected heterogeneity among mesodermal and ectodermal cells. The new scRNA-seq analysis identified a previously undescribed subpopulation of cells expressing *abcc9*, a marker of mammalian pericytes, a perivascular cell type of mesodermal origin [46] (Figure 5B). Interestingly, *abcc9* is also expressed in the larval mesoderm of *C. teleta* (Figure 6D–F), suggesting an evolutionarily conserved function of this gene in mesodermal differentiation. Likewise, the new dataset captured four transcriptionally distinct ectodermal subpopulations: *nox5*^+^/*y325*^+^ cells, *rsph1*^+^/tubulin^high^ cells, *foxred2*^+^ cells, and *zonadhesin*^+^ cells. Whole-mount *in situ* hybridisation of *foxred2*, *rsph1* and *zonadhesin* confirmed their ectodermal expression (Figure 6G–O). The flavoprotein *foxred2* [47] (Figure 6G–I) and the axonemal component *rsph1* [48] (Figure 6J–L) are broadly expressed in the ectoderm, albeit with distinct patterns. However, *zonadhesin*, an adhesion protein that in humans localises to the sperm acrosome and binds to the zona pellucida of the egg [49], is expressed in a subpopulation of large ectodermal cells (or clusters of cells), mainly concentrated in the anterior and posterior ends of the larvae (Figure 6M–O). The new chromosome-level assembly and updated annotations, therefore, enhance cluster resolution and cell-type identification in scRNA-seq analyses, supporting the identification of previously overlooked cell types and cell-type-specific markers that can be the subject of future study.

### A new genome browser for C. teleta

To facilitate the use of the genomic data generated during this study, we have developed the Capitella Genome Project Portal, located at (weblink). This portal serves as a centralised resource for genome annotations and additional biological information, including ATAC-seq and mitochondrial genome data. Additionally, the portal features an interactive JBrowse [50] front-end that allows users to visualise genomic data, as well as a custom interface for performing BLAST queries on these data [51].

## Conclusion

Our study presents the chromosome-level assembly of the laboratory strain of the polychaete worm *C. teleta*, a well-established invertebrate research system in evolutionary and developmental biology, comparative epigenomics, ecotoxicology, and conservation [7, 11, 52]. By optimising long-read sequencing and chromatin conformation capture technologies in this annelid, we produced a reference genome assembly that matches its karyotype and expected genome length, removes haplotypic retention, and contains an improved annotation of its repeat landscape and protein-coding gene repertoire. This new genome assembly confirms that *C. teleta* has a slow-evolving gene repertoire, retaining a significant fraction of the ancestral metazoan gene families. Paradoxically, the conservative evolutionary dynamics at the molecular and gene level are not reflected at the chromosomal level, as *C. teleta* displays heavily rearranged nuclear and mitochondrial genomes. As anticipated, a chromosome-level assembly enhances the analysis of functional genomic data by increasing the mappability of RNA-seq, ATAC-seq, and EM-seq data, facilitating the detection of gene expression, peak calling, and DNA methylation profiling. This is better illustrated by the analysis of scRNA-seq data, where the new annotation resolves more cells and clusters, identifying new cell-type-specific markers and cellular populations. Overall, this reference genome addresses a critical community need, bringing the molecular study of *C. teleta* to current gold standards comparable to those of other annelid research organisms, such as *O. fusiformis* and *Platynereis dumerilii* [18, 53].

In addition to the new chromosome-level assembly, this study also establishes a publicly available online genome browser that integrates the new sequencing data with existing functional genomic resources. This dedicated online platform ensures that the available genomic resources for *C. teleta* comply with FAIR principles, making them easily findable, accessible, interoperable, and reusable. We anticipate that this will have a lasting impact on the research community investigating *C. teleta* and other invertebrate species, facilitating their access to multimodal and comparative genomic datasets that will enrich and enable new research avenues, from the implementation of cutting-edge epigenomic profiling approaches and genome editing to phylogenomic and environmental molecular monitoring.

## Methods

### Animal culture and collections

We followed established protocols to maintain a stable colony of *C. teleta* Blake, Grassle & Eckelbarger, 2009 [5]. For genomic DNA extractions, adult worms were collected and washed in artificial seawater (ASW) before snap-freezing in liquid nitrogen. For gene expression analyses, brood tubes were dissected to collect larvae at stages 5 and 7, which were anaesthetised in MgCl_2_ in ASW for 10 minutes before fixation in 4% paraformaldehyde (ThermoFisher) in ASW overnight at 4 °C. After washing out the fixative in PBS + 0.1% Tween-20, the samples were gradually dehydrated in 100% MeOH and eventually stored in pure 100% methanol at –20 °C for downstream staining protocols.

### Genomic DNA extractions

High molecular weight (HMW) gDNA was extracted from adult worms using the MagAttract HMW DNA Kit (Qiagen, 67563) with slight modifications (Supplementary Table 1). Briefly, 100 mg of tissue were washed three times in Phosphate Buffer Saline (PBS) until the solution was clear, and tissue lysis was then conducted in Buffer ATL supplemented with proteinase K at 56 °C, shaking at 900 rpm for 15-20 minutes. The remaining steps were performed according to the manufacturer’s recommendations.

### Genome sequencing, assembly, and quality check

Nanopore sequencing was performed in r9.4.1 and r10.4.1 pores (Supplementary Table 1), and the resulting reads were base-called with Guppy (https://nanoporetech.com/software/other/guppy) using the super accurate (SUP) mode, producing a total of 103 Gb from passed reads and an N50 of 1.2 Kb. All passed reads were then merged and filtered with Nanofilt (v.2.8.0) [54] to select reads greater than 3.8kb.

Nanopore reads were assembled using Canu (v2.2) [55] using “batOptions= -dg 3 -db 3 -dr 1 -ca 500 -cp 50”. The genome was then polished based on long reads greater than 1kb and short Illumina reads using HyPo (v.1.0.3) [56]. Purge Haplotigs (v.1.1.2) [57] with parameters (-l 65, -m 145, -h 300) was used to reconstruct a high-quality haploid reference assembly, which was further decontaminated following the BlobTools (v1.1) pipeline [58], removing contigs with a non-animal taxonomy. BUSCO (v.5.4.4) and QUAST (v.4.6.1) were used to assess genome completeness and evaluate assembly quality [59, 60].

### Chromosome-scale scaffolding

Hi-C reads were generated from a single individual using Proximo Hi-C (Phase Genomics), resulting in ∼60M reads. Reads were mapped following guidelines from Arima Genomics (https://github.com/ArimaGenomics/mapping_pipeline). Reads were first mapped to the assembly with BWA-MEM (v.0.7.17) as single reads. The 3’-end of reads comprising a ligation junction was then filtered using ‘filter_five_end.pl’ before combining read pairs with ‘two_read_bam_combiner.pl’. Assembly scaffolding was performed using YaHS (v.1.2a) [61], resulting in 11 chromosome-level scaffolds corresponding to 10 chromosomal units.

### Annotation of repeats and transposable elements

Repeat elements were annotated with RepeatModeler (v.2.0.5) [62] and RepBase, constructing a *de novo* repeat library for *C. teleta*. After potentially *bona fide* genes were removed from the repeat library, repeat elements were annotated with RepeatMasker (v.4.1.5) (http://repeatmasker.org). To estimate Kimura substitution levels, we used the scripts “calcDivergenceFromAlign.pl” and a custom-modified version of “createRepeatLandscape.pl” from RepeatMasker v4.1.0.

### Gene prediction and functional annotation

Protein-coding genes were annotated following a previously established pipeline [18]. Briefly, all available RNA-seq datasets were mapped to the soft-masked genome using STAR (v.2.7.11b) [63], and a curated set of splice junctions was created with Portcullis (v.1.2.4) [64]. The Mikado (v.2.3.4) pipeline [65] was used to merge all transcriptomic evidence into a single set of best-supported gene models. These were then used to train Augustus (v.3.5.0) [66, 67] and, together with the splice junctions from Portcullis and the spliced alignment of the *O. fusiformis* proteome on *C. teleta* genome with Miniprot (v0.8-r220) [68], generate exon, intron, and gene hints, respectively, for *ab initio* gene prediction with Augustus (v.3.5.0). Finally, the PASA (v.2.5.2) pipeline [69] was used to combine RNA-seq and *ab initio* gene models into a final gene set. From this, we removed spurious predictions with in-frame stop codons, predictions that overlapped with repeats, and those that had high similarity to TEs in the RepeatPeps.lib database. This filtered gene set comprised 26,591 genes and 34,679 different transcripts. To assess the completeness of this annotation, we ran BUSCO (v.5.4.4) [59] in proteome mode, which yielded 95.7% of the core genes present. The Trinotate pipeline (v.4.0.0) [70] and PANTHER (v.2.2) [71] were used to provide functional annotations for the filtered gene models. For assembly comparisons, output gff annotations from the previous and new genome assemblies were mined in a Python script, with AGAT [72] used to extract intronic regions.

### Gene family evolution analyses

Non-redundant proteomes with the longest isoform per gene were used to calculate orthogroups (i.e., gene families) using OrthoFinder (v.2.5.5) [73], MMseqs2 [74], and IQ-TREE reconstruction [75]. These gene families were parsed and mapped onto a reference species phylogeny to infer gene family gains and losses at different nodes and tips using the ETEL3 library [76], and to estimate the node of origin for each gene family. Gene expansions were computed for each species using a hypergeometric test against the median gene number per species within a given family, as previously published [27]. Gene Orthology (GO) enrichment analysis of lost orthogroups (using the functional annotation of the respective genes in *O. fusiformis*) and expanded gene families was performed using the topGO (v.2.60.1) package [77] and a Fisher’s exact test.

### Macrosynteny analyses

Chromosome-level assemblies for *C. teleta*, *U. unicinctus*, *P. maximus*, and *L. longissimus* were annotated with MetaEuk (v.6-a5d39d9) [78], and the rbhXpress (v.1.2.3) script was used to identify reciprocal best BLAST hit homologues that were plotted as Oxford grid plots and chord diagrams with the macrosyntR package (v.0.2.19) [79]. Ancestral linkage groups were assigned based on inter-species chromosomal correspondences using as a reference the chromosomal annotations of *P. maximus*, *U. unicinctus* and *L. longissimus* from previous studies [24].

### Assembly and annotation of the mitochondrial genome

The complete mitochondrial genome of *C. teleta* was obtained by BLASTN sequence similarity searches of the complete mitochondrial genome of *Notomastus sp.* (NCBI LC661358.1) against a *de novo* assembled transcriptome [18]. The genome was preliminarily annotated using the MITOS 2 web server [80] with default settings and the invertebrate mitochondrial genetic code (option 5). Protein-coding genes were edited manually based on the annotation of Open Reading Frame Finder (https://www.ncbi.nlm.nih.gov/orffinder/). A circle plot displaying mitochondrial gene order, GC content, and GC skew was generated using the online tool CGView [81], and the resulting SVG file was edited in Adobe Illustrator. To validate the mitochondrial gene order, six primer pairs were designed to amplify regions overlapping the boundaries between neighbouring genes. PCR products were purified using a Monarch PCR DNA cleanup kit (New England Biolabs, #T1130) and Sanger sequenced. The resulting sequences were aligned to the original mitochondrial genome assembly using ApE [82], confirming mitochondrial gene order.

### cox1 phylogeny of the genus Capitella

A phylogeny of the mitochondrial *cox1* gene was produced for the genus *Capitella*, with *Notomastus* as the outgroup. An initial BLAST search of NCBI databases was performed against the *C. teleta cox1* gene, filtering for taxa within the *Capitella* genus, yielding a dataset of 291 *cox1* sequences (mostly partial CDS). In addition, the *cox1* sequence from our lab strain of *C. teleta*, the designated species holotype of *C. teleta* [5], and the *cox1* sequence of *Notomastus sp.* were included in the final dataset. Sequences were aligned via the online tool Clustal Omega [83], trimmed using TrimAl (v1.5.0) [84], and a maximum-likelihood analysis of the aligned dataset was performed using IQ-TREE2 (v2.3.6) [75], applying an ultrafast bootstrap of 5000 replications [85]. The resulting consensus tree was viewed and edited using FigTree v1.4.4 (https://github.com/rambaut/figtree) and manually rooted to *Notomastus* as the chosen outgroup. The tree was then annotated and further edited in Adobe Illustrator.

### Bulk transcriptomic profiling

Batch correction was performed using the SVA package (v. 3.46.0) [86], which employs surrogate variable analysis (SVA) to identify groupings of hidden and unwanted variation. This was achieved by creating a null model that contained the known variables and a full model that included the added variable of interest. The SVA function then estimates and removes surrogate variables. Limma (v.3.54.2) [87] and DESeq2 (v.1.38.3) [88] were then used to remove batch effects and normalise gene expression levels. Transcripts were then clustered according to their normalised DESeq2 and batch corrected expression dynamics through soft *k*-means clustering using the Mfuzz (v.2.58.0) package [89] using an optimal number of 10 clusters (inferred through the elbow method to the minimum centroid distance as a function of the number of clusters). Transcripts with no expression at any developmental stage (346 out of 34679) were discarded.

### ATAC-seq profiling

We used cutadapt (v.2.5) [90] to remove sequencing adaptors and trim paired 150-base reads to 75 bases. Quality-filtered reads were mapped using NextGenMap (v.0.5.5) [91] in paired-end mode, duplicates were removed using SAMtools (v.1.9) [92], and mapped reads were shifted using deepTools (v.3.4.3) [93]. Peak calling was performed using MACS3 (v.3.0.3) [94] with the following settings: -f BAMPE --min-length 100 --max-gap 75 and -q 0.01.

Peaks in repetitive regions were filtered using BEDTools (v.2.31.1) [95]. Reproducible peaks were identified using IDR (Irreproducible Discovery Rate) [96] with flag --soft-idr-threshold 0.05. DiffBind (v.3.14.0) [97] was used to generate and normalise the final consensus peak set and pairwise contrast of peaks between two stages. Peak clustering according to accessibility dynamics was performed using the Mfuzz (v.2.64.0) package [89]. Principal component analyses were performed on the variance-stabilising-transformed DiffBind-normalised matrices. Peak annotation was performed using HOMER (v.5.1) [98]. The Pearson correlation coefficient between chromatin accessibility and gene expression was computed individually for each peak using two-sided tests.

### EM-seq profiling

Following the library preparation protocol described in Guynes et al. [16], adaptors and low-quality reads were trimmed using Trim Galore! (v0.6.7) [90] with the parameters -j 8 -e 0.1 -q 20 -O 1. Cleaned reads were then aligned to the genome using Bismark (v0.24.0) [99] with the options --q --local --score-min G,20,8 --ignore-quals --no-mixed --no-discordant -- dovetail --maxins 500. Methylation was called for all cytosine contexts using the bismark_methylation_extractor tool, and the resulting coverage files were used for all downstream methylation analyses with custom R scripts [16].

### Single-cell RNA-seq analyses

For scRNA-seq analysis, we downloaded previously published scRNA-seq reads for *C. teleta* from the Sequence Read Archive (BioProject PRJNA669754) [20]. Alignment of these previously obtained sequencing reads and processing into a digital gene expression matrix were performed using 10x Genomics Cell Ranger version 8.0.0 [100], with STAR version 2.7.2a [63] as the aligner, using standard parameters. Cell Ranger was run separately with the same set of reads and parameters for the old and new *C. teleta* genome. Single-cell transcriptomes were then processed and analysed using Seurat version 5.1.0 [101] and R version 4.4.1 [102]. Cells that had a low (≤300 genes and UMIs detected) or high number of detected features and UMIs (≥1500 genes and UMIs detected) were removed. They were then log-normalised (Seurat:NormalizeData, scale.factor = 10000) and scaled. For visualisation, we identified the top 2000 variable genes (Seurat::FindVariableFeatures, selection.method = “vst”, nfeatures = 2000), performed PCA (Seurat::RunPCA), and identified significant PCs (Seurat::JackStraw, dims=100). A Uniform Manifold Approximation and Projection (UMAP) was calculated using 25 nearest neighbours and the 14 most significant PCs (Seurat::RunUMAP, n.neighbors = 25). For clustering, we used the Leiden approach on the top 14 PCs (Seurat::FindClusters, algorithm = 4, resolution = c(0.5,0.75,1,2,3), n.start = 50, random.seed = 17). Markers were calculated for roughly annotated clusters at each resolution. The different resolutions were assessed manually, and a resolution was chosen based on the clustering that seemed most biologically relevant. For both datasets (aligned to old and new genome), at higher resolutions, subclusters without sufficient differentially expressed genes were combined to ensure that each cluster was sufficiently distinct – marker genes for individual clusters were identified using ROC and Wilcoxon Rank Sum Tests using the command Seurat:FindMarkers(test.use = “roc”/“wilcox”, min.pct = 0.25, logfc.threshold = 0.25). Subclusters without at least two differentially expressed genes were merged. Such manual curation was performed to represent the comparative molecular heterogeneity of cell types recovered in our data when aligned to the old and new *C. teleta* genomes.

### Gene expression analyses

Riboprobes of ∼1–1.5 kb for candidate cell-type-specific markers were synthesised as previously described [103] and validated by Sanger sequencing. Colourimetric whole-mount *in situ* hybridisation was performed using an established protocol [39], where samples were incubated with 1.5–2 ng/μL of riboprobes at 62 °C for 72 hours. After probe washes, samples were incubated overnight at 4 °C with anti-DIG-AP Fab fragments (Sigma) diluted 1:4500. Signal development with NBT and BCIP (Sigma) was done at 27 °C and checked regularly.

### Imaging

Representative larvae from whole-mount *in situ* hybridisation analyses were imaged using a Leica DMRA2 upright microscope equipped with an Infinity5 camera (Lumenera), employing differential interference contrast optics. Focus-stacking software (Helicon Focus Lite) was used to combine a stack into a single, entirely focused image. Colour balance was adjusted in Photoshop (Adobe), applying changes to the entire image and not just parts of it. Final figures were designed using Illustrator (Adobe).

## Data availability

The newly generated data from this whole-genome sequencing project have been deposited in DDBJ/ENA/GenBank under the project accession PRJEB101787. The *Capitella* Genome Project portal (weblink) offers a comprehensive resource for this species, featuring a BLAST interface, a genome browser, DNA and protein sequence downloads, functional annotation of gene models, and ATAC-seq and EM-seq data.

## Author contributions

ECS, ADB, and JMMD designed and conceived the study. BED, ECS, and PG conducted data generation. BED, PG, and JMMD conducted genome assembly and annotation. BED, JW, and JMMD performed genome assembly comparisons, gene family evolutionary analyses, and macrosynteny reconstruction. TF and JW assembled, annotated, and analysed the mitochondrial genome. BED and RDD performed bulk transcriptomic analyses. YL and JW analysed ATAC-seq data, and KG analysed EM-seq data. AS performed scRNA-seq analyses, and AMCB and JM performed gene expression validation. RTM, SS, SZ, RY, TGW, and ADB conceptualised and developed the *Capitella* Genome Project Portal. All authors contributed to drafting the manuscript.

## Acknowledgements

We thank past and present members of the Martín-Durán lab for their help and contributions to establishing *C. teleta* in the lab, as well as the technical support staff at the Department of Biology at Queen Mary University of London. We also thank Philipp Rescheneder, Sean McKenzie, and Vania Costa from Oxford Nanopore Technologies for their assistance with long-read sequencing in *C. teleta*. This study utilised computational resources from the High-Performance Computing Facility Apocrita at Queen Mary University of London. This work was funded by grants to JMM-D from the European Union Horizon 2020 Framework Programme under the European Research Council (ERC Starting Grant agreement number 801669), the Biotechnology and Biological Sciences Research Council (BB/T008709/1 and BB/Y004221/1) and the Leverhulme Trust (RPG-2023-121), as well as the China Scholarship Council doctoral fellowship to JW (CSC NO. 202306330025). This research was supported in part by the Intramural Research Program of the National Institutes of Health [ZIA HG000140 to ADB]. The contributions of the NIH authors are considered Works of the United States Government. The findings and conclusions presented in this paper are those of the authors and do not necessarily reflect the views of the NIH or the U.S. Department of Health and Human Services.

---

**Supplementary Figure 1.**
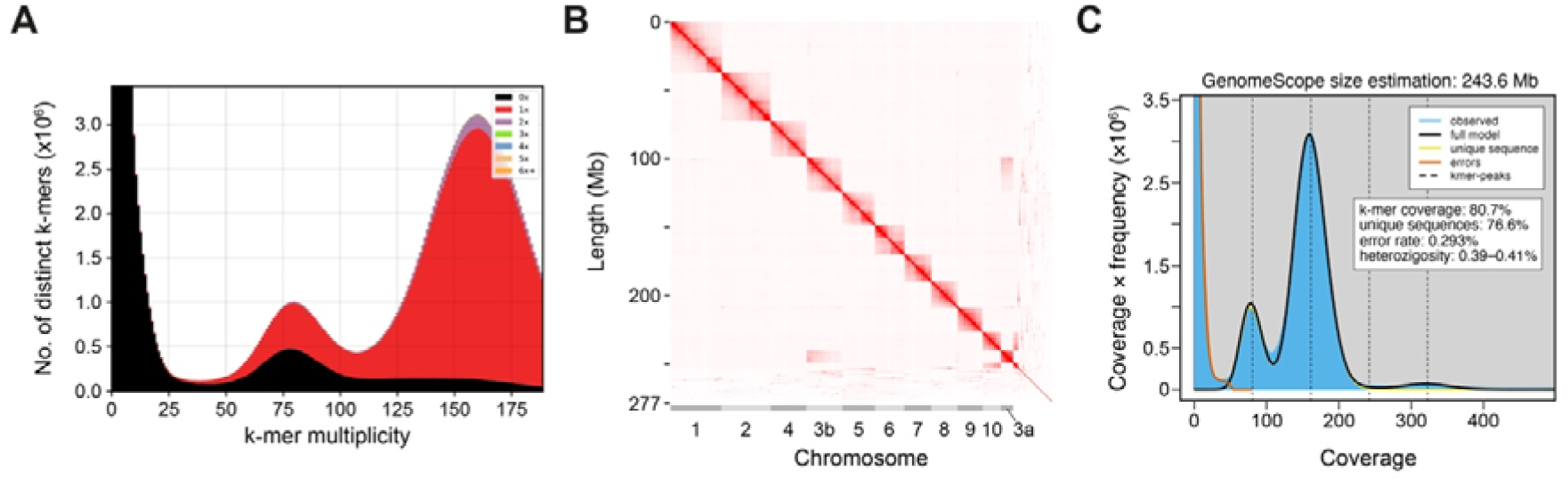
Assembly and scaffolding of *C. teleta*’s genome. (**A**) *k*-mer distribution plot after dehaploidisation. The final assembly primarily consists of a single haplotype. (**B**) HiC contact heatmap showing the scaffolding of the initial assembly into 10 pseudo-chromosome molecules, with chromosome 3 not fully resolved and split into two (3a and 3b). (**C**) *k*-mer distribution plot estimates *C. teleta*’s genome size in 243.6 Mb and a low heterozygosity level of 0.39–0.41%, as expected for an inbred cultured line.

**Supplementary Figure 2.**
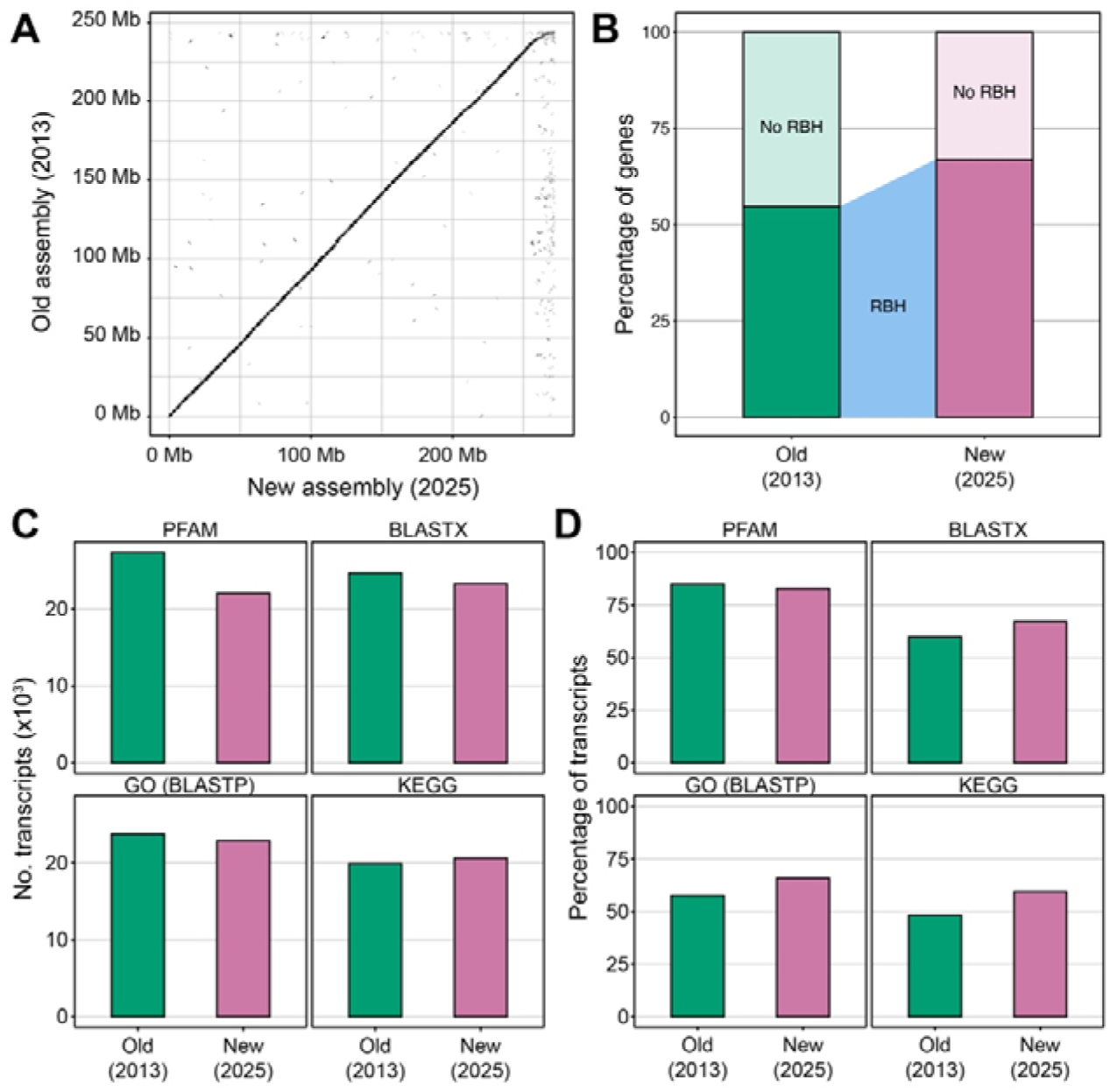
Quality and completeness comparison between the old (2013) and new (2025) genome assemblies and their respective gene annotations. (**A**) A whole-genome dot plot comparing the old and new assemblies reveals high similarity. (**B**) The percentage of genes that have a Reciprocal Best Hit (RBH) between the old and new assemblies is substantially higher in the new assembly (66.93%) than in the old (54.61%), indicating a less redundant gene set. (**C**) The absolute number of transcripts with functional annotations from PFAM, BLASTX, GO, and KEGG databases is slightly lower for the new gene set. (**D**) In contrast, the percentage of transcripts with functional annotations is significantly higher in the new gene set. Together, (**B**–**D**) show that the new annotation produced a higher-quality set of gene models with fewer spurious and redundant gene predictions.

**Supplementary Figure 3.**
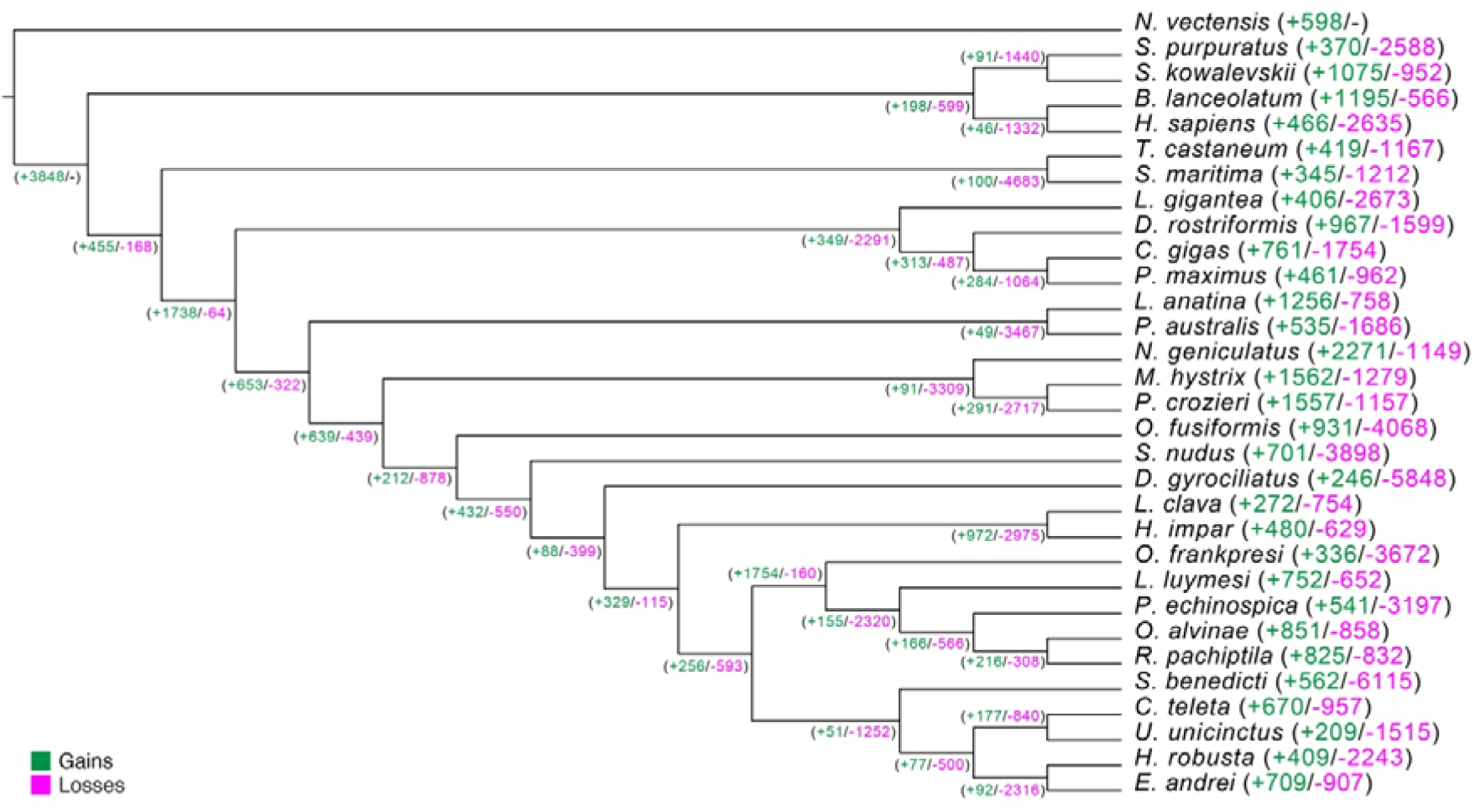
Detailed cladogram of gene family evolution across Metazoa. Inferred history of gene family gains and losses for all species included in this study, using a consensus tree topology of Bilateria, Spiralia and Annelida [3]. The numbers at each internal node and along each terminal branch correspond to the number of gene families inferred to have been gained (+) and lost (-) on that lineage/node. Colours distinguish gains (green) from losses (magenta).

**Supplementary Figure 4.**
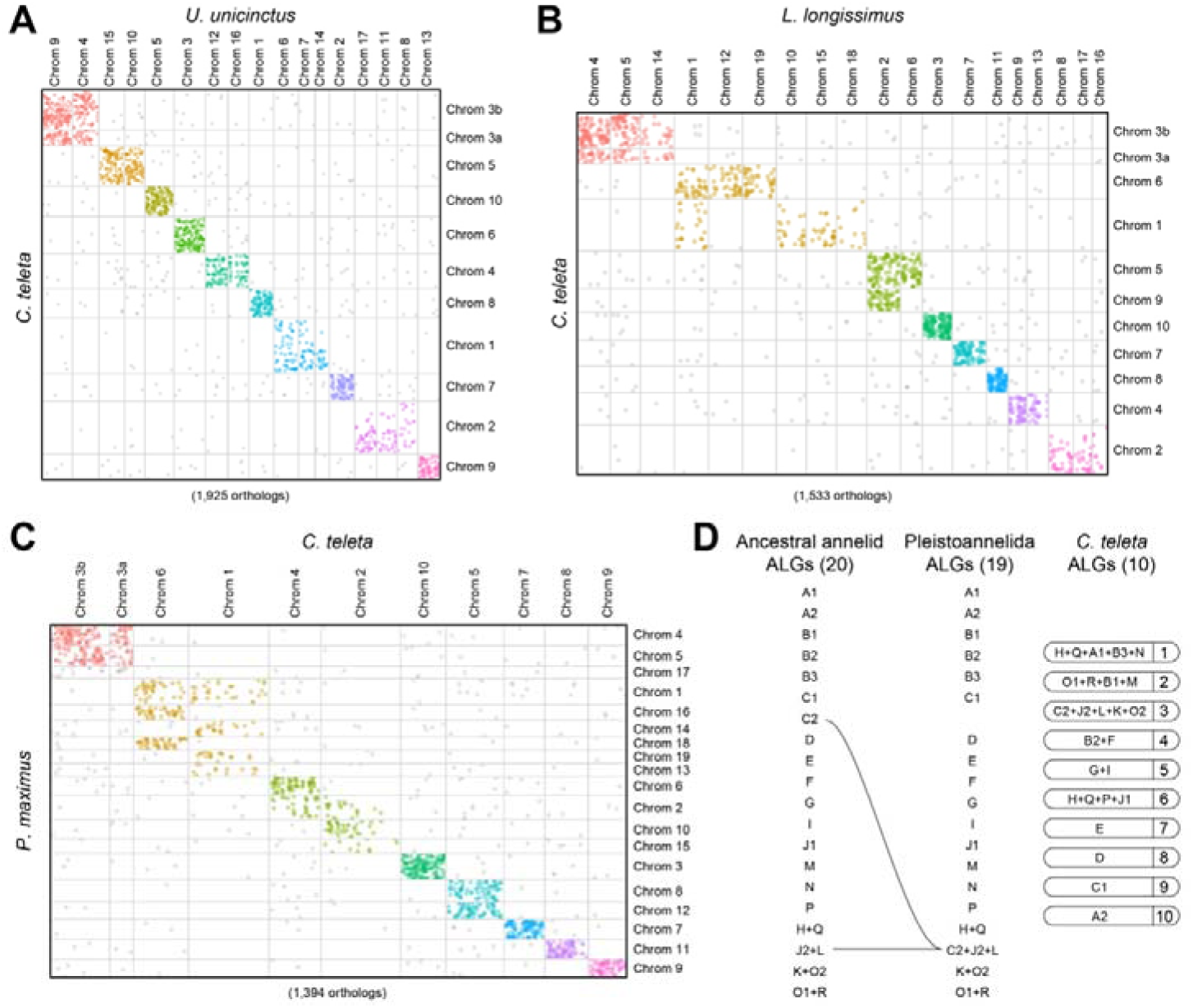
Macrosynteny and a proposed model of chromosome evolution in Annelida. (A, **B)** Dot plots comparing the chromosome-level assembly of *Capitella teleta* against the annelids *Urechis unicinctus* (A) and *Lineus longissimus* (B). The strong diagonal patterns, where each colored block represents a pair of homologous chromosomes, indicate a high degree of chromosome structure conservation within Annelida. **(C)** A macrosynteny comparison between *C. teleta* (Annelida) and the scallop *Pecten maximus* (Mollusca). While one-to-one chromosome homology is lost between phyla, the persistence of numerous conserved syntenic blocks points to a shared ancestral spiralian karyotype. **(D)** A schematic model for the evolution of the *C. teleta* karyotype (n=10). This model proposes a specific series of chromosome fusion events, tracing the lineage from a putative ancestral annelid with 20 ancestral linkage groups (ALGs) through an intermediate Pleistoannelida ancestor.

**Supplementary Figure 5.**
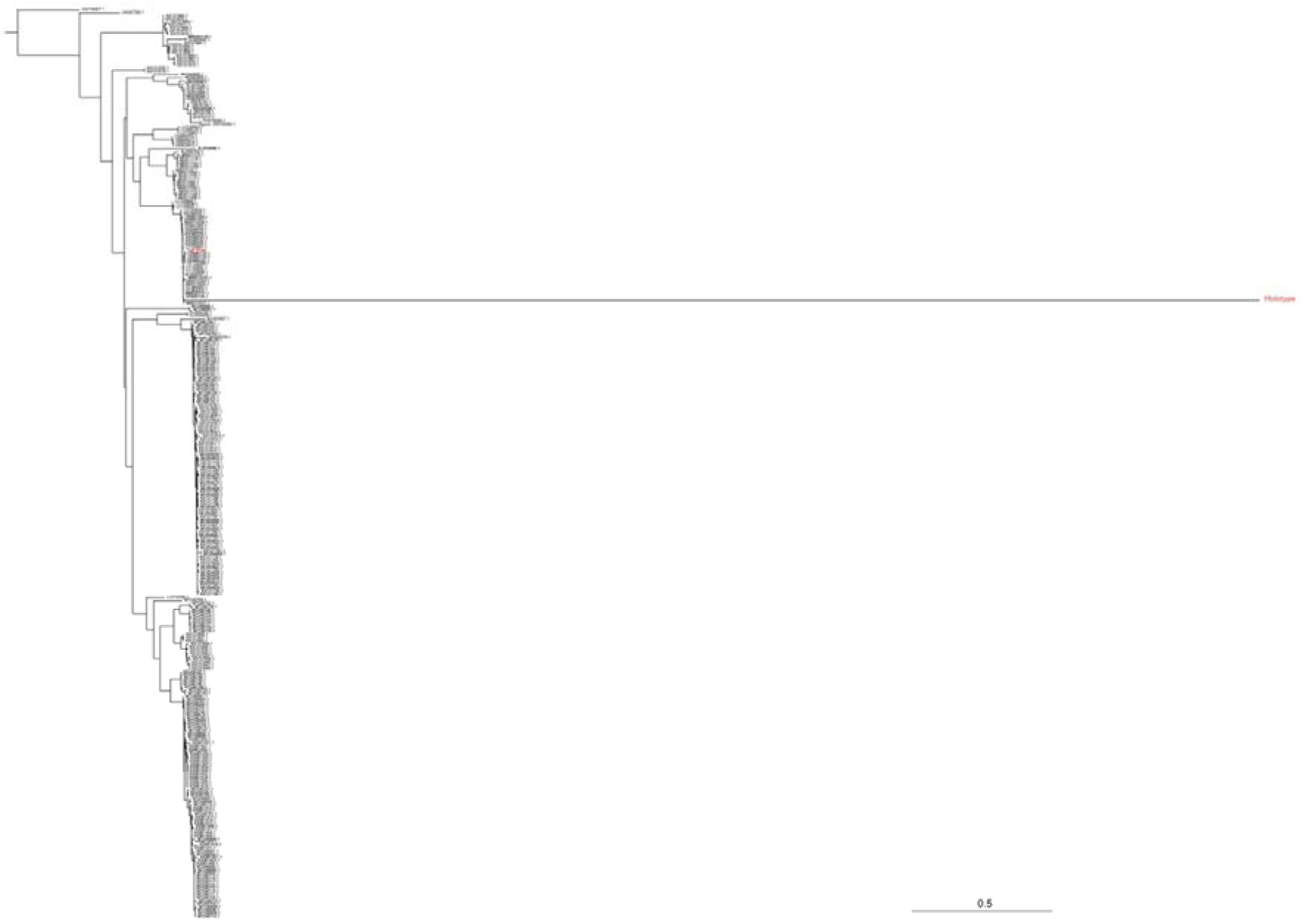
The phylogeny of the genus *Capitella*. Phylogenetic tree reconstructed with IQ-Tree using the available sequences for the mitochondrial *cox1* gene. The sequences from the holotype and this study are in red. Note how the holotype sequence is dramatically divergent compared with the others.

**Supplementary Figure 6.**
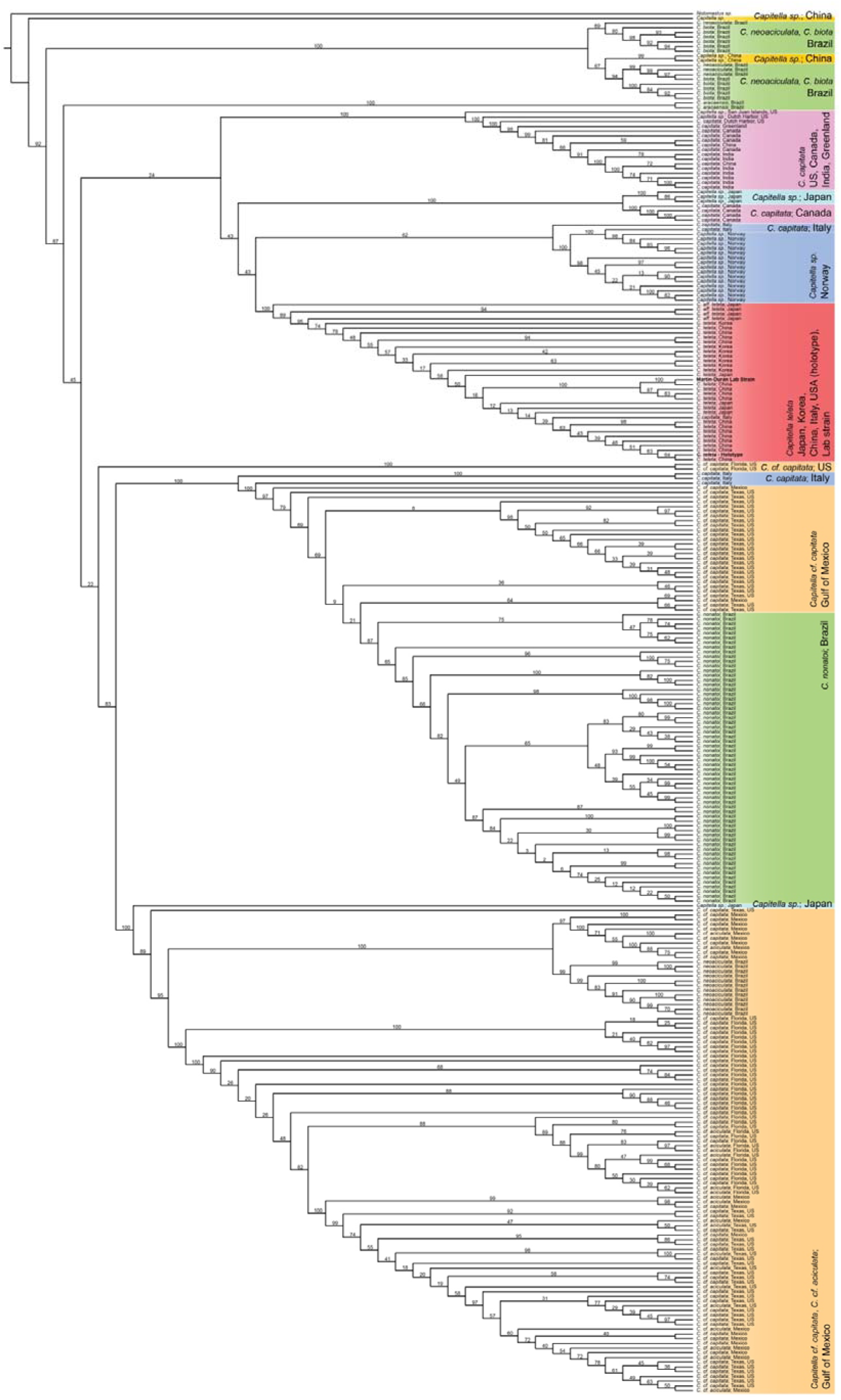
The phylogenetic relationship and geographical distribution of the genus *Capitella*. Cladogram indicating the phylogenetic relationships of the different species within *Capitella* based on the available *cox1* mitochondrial gene sequences. The tree topology is based on Supplementary Figure 5. The coloured boxes indicate major clades based on geographical distribution and assumed species identification. Note, however, that some species, like *C. capitata* and *C. neoaciculata*, are not monophyletic clades and may, thus, represent cryptic species.

**Supplementary Figure 7.**
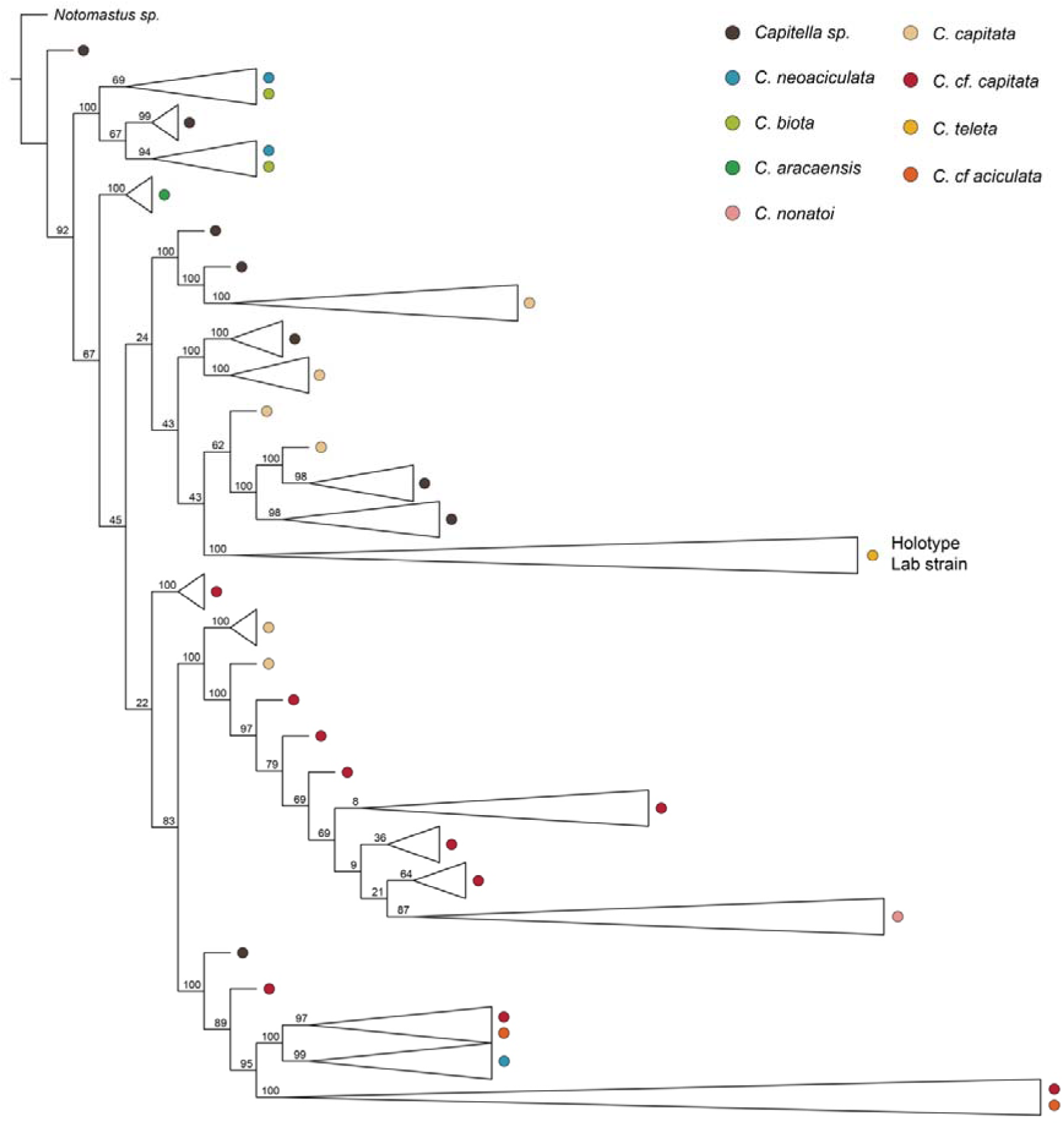
The phylogenetic relationships of the genus *Capitella*. Cladogram indicating the main groups obtained through phylogenetic reconstruction using the *cox1* mitochondrial gene. The tree topology is based on Supplementary Figure 5. The coloured circles indicate the species included in the collapsed clades.

**Supplementary Figure 8.**
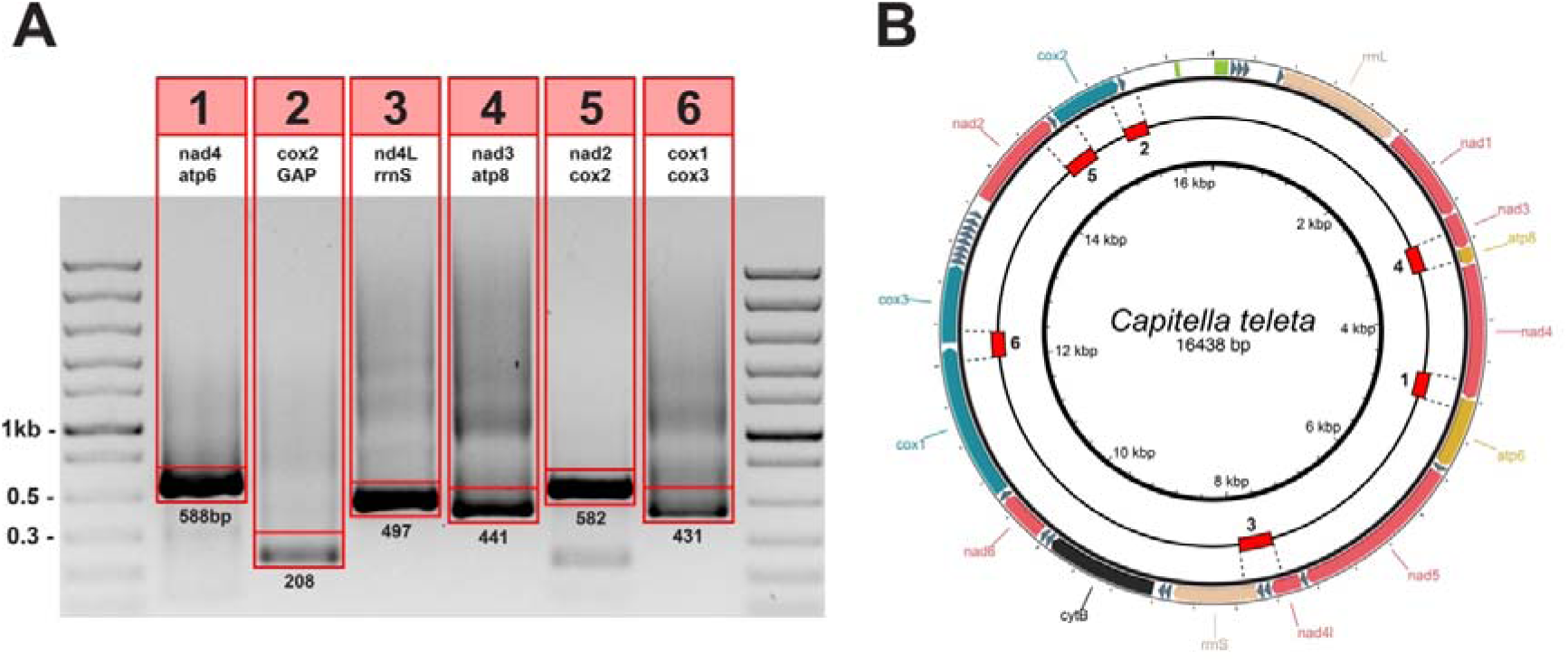
Experimental validation of the mitochondrial genome assembly in *C. teleta*. (**A**) Diagnostic PCR of six amplicons located at positions in the mitochondrial genome assembly of *C. teleta* (shown as red boxes in **B**) that correspond to syntenic rearrangements.

**Supplementary Figure 9.**
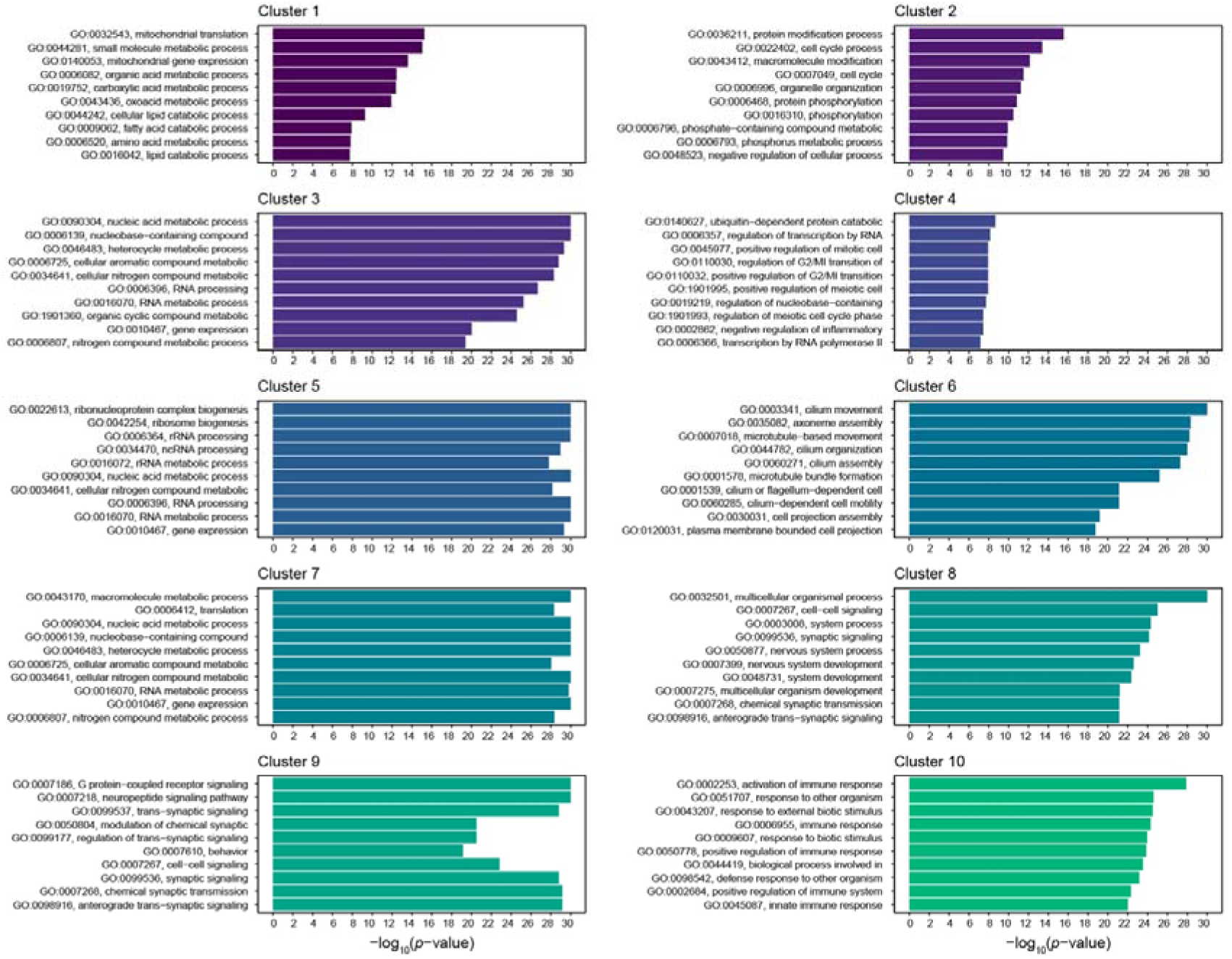
Gene Ontology enrichment in clusters of temporally coregulated genes. Bar plots depicting the top 10 Gene Ontology (GO) terms in each cluster of temporally coregulated genes obtained after soft *k*-means clustering of the time course of bulk RNA-seq data.

**Supplementary Figure 10.**
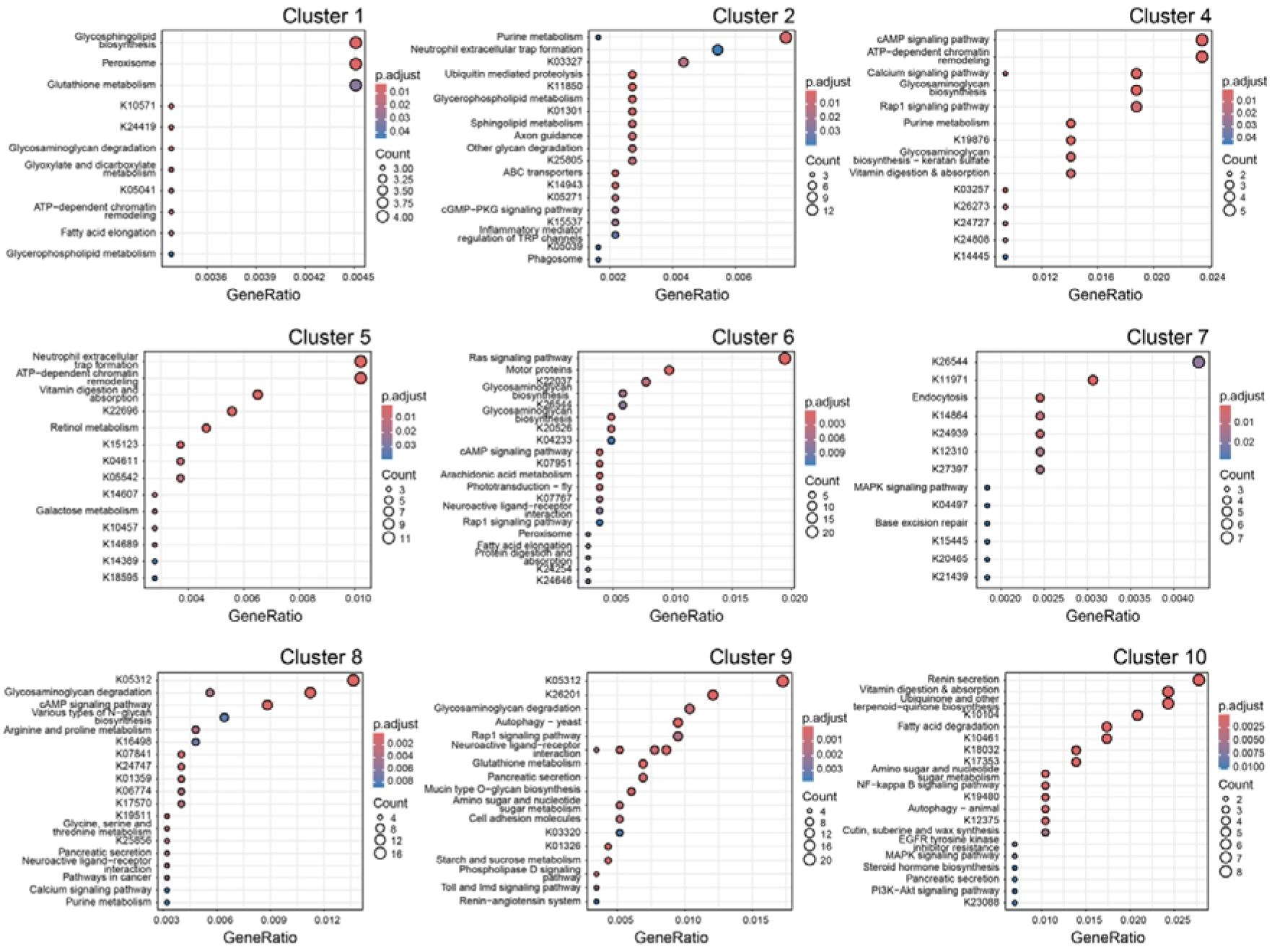
KEGG enrichment in clusters of temporally coregulated genes. Dot plots depicting the differentially enriched KEGG terms in each cluster of temporally coregulated genes obtained after soft *k*-means clustering of the time course of bulk RNA-seq data. There were no differentially enriched KEGG terms in Cluster 3.

**Supplementary Figure 11.**
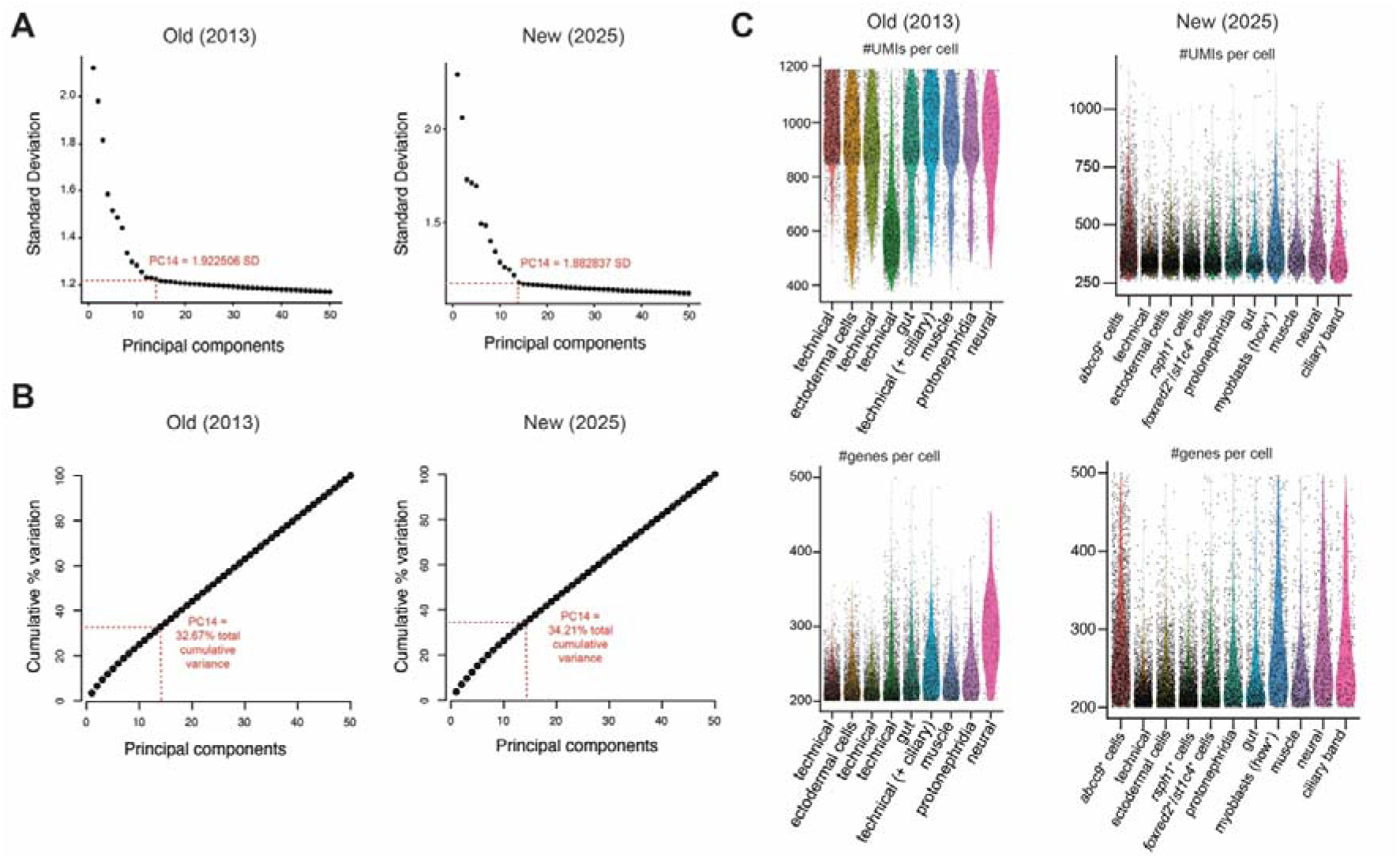
Standard variance and principal component comparisons for *C. teleta* scRNA-seq analysis between old and new annotations. (**A**) Elbow plots showing standard deviation plotted against the number of principal components (PCs) using whole-larvae *C. teleta* scRNA-seq data quantified using the indicated annotation. Standard deviation values for 50 PCs are shown in both cases. (**B**) Plots of cumulative percentage variation against the number of PCs from the scRNA-seq data quantified using the indicated annotation. Total cumulative variance values for 50 PCs are shown for each annotation. (**C**) Violin Plot showing the number of UMIs and genes per cluster between the old and new annotations as indicated. No clusters have zero genes or UMIs.

**Supplementary Figure 12.**
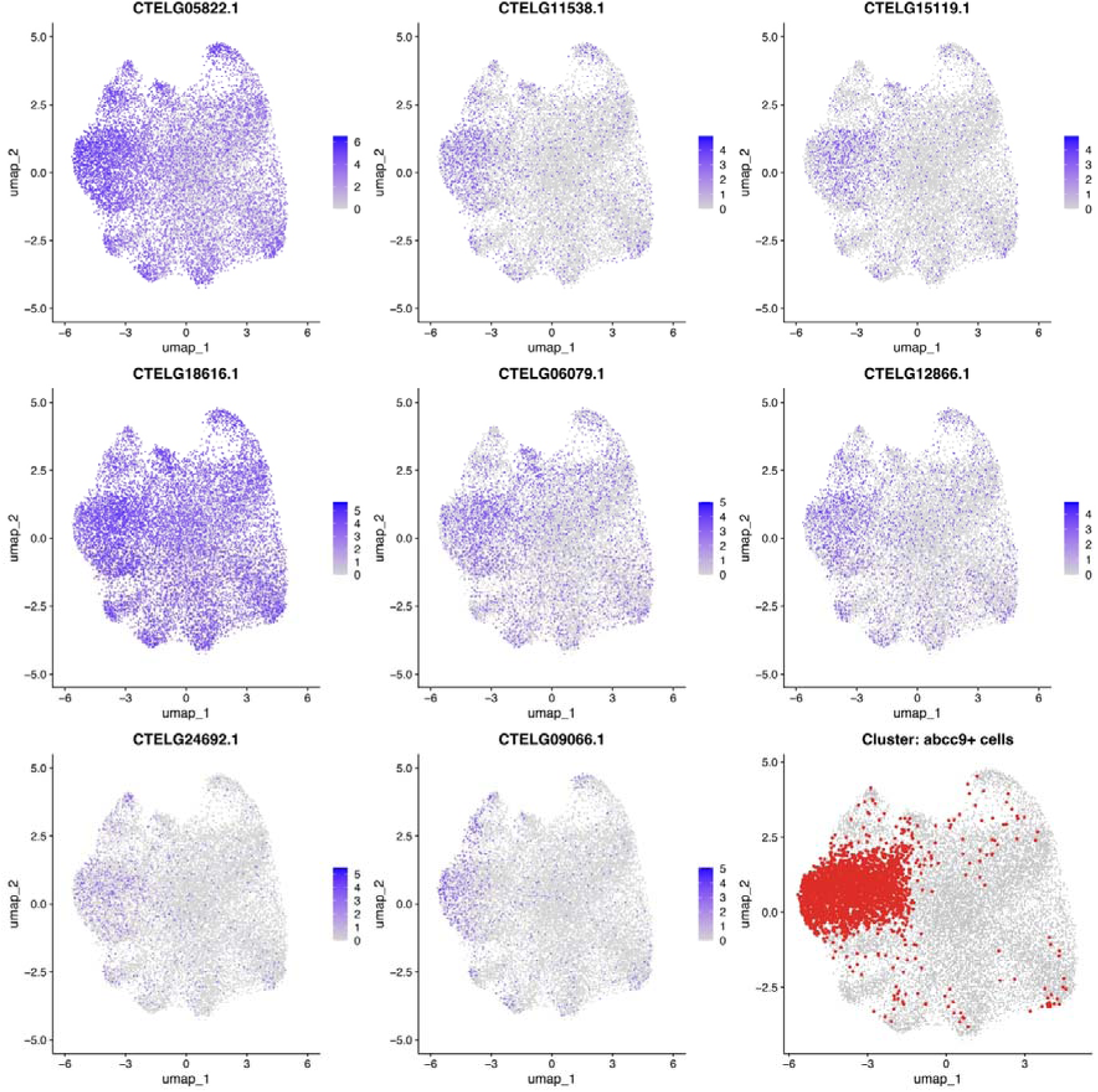
UMAP plots for the top eight marker genes for cluster 1 (*abcc9*^+^ mesoderm cells). UMAP plots are coloured based on expression levels obtained from the Wilcoxon Rank Sum differential expression test. The colour bar indicates the expression level (natural log) of each gene.

**Supplementary Figure 13.**
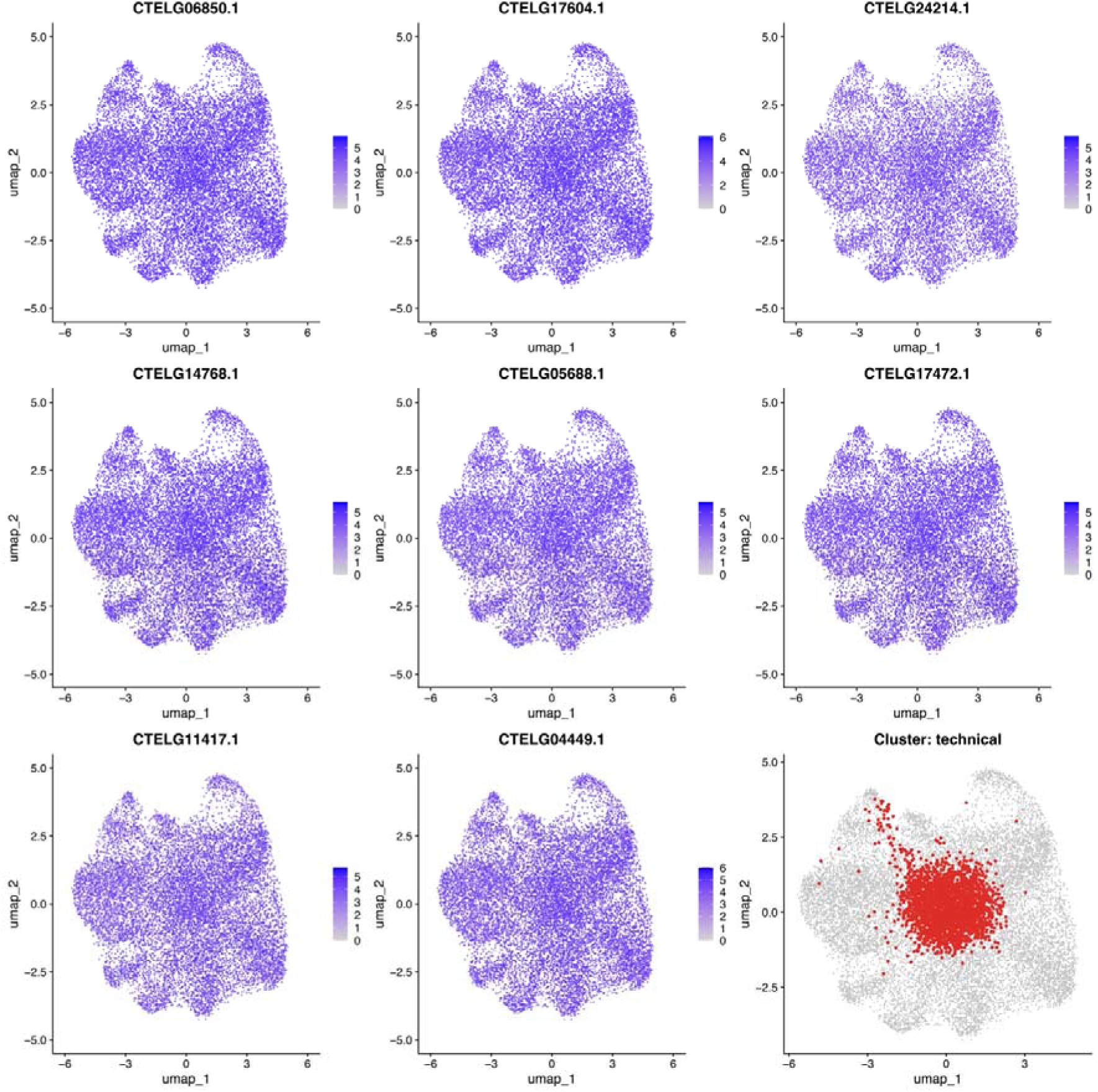
UMAP plots for the top eight marker genes for cluster 2 (technical). UMAP plots are coloured based on expression levels obtained from the Wilcoxon Rank Sum differential expression test. The colour bar indicates the expression level (natural log) of each gene. Note the ubiquitous expression of these genes, indicating a “technical” cluster.

**Supplementary Figure 14.**
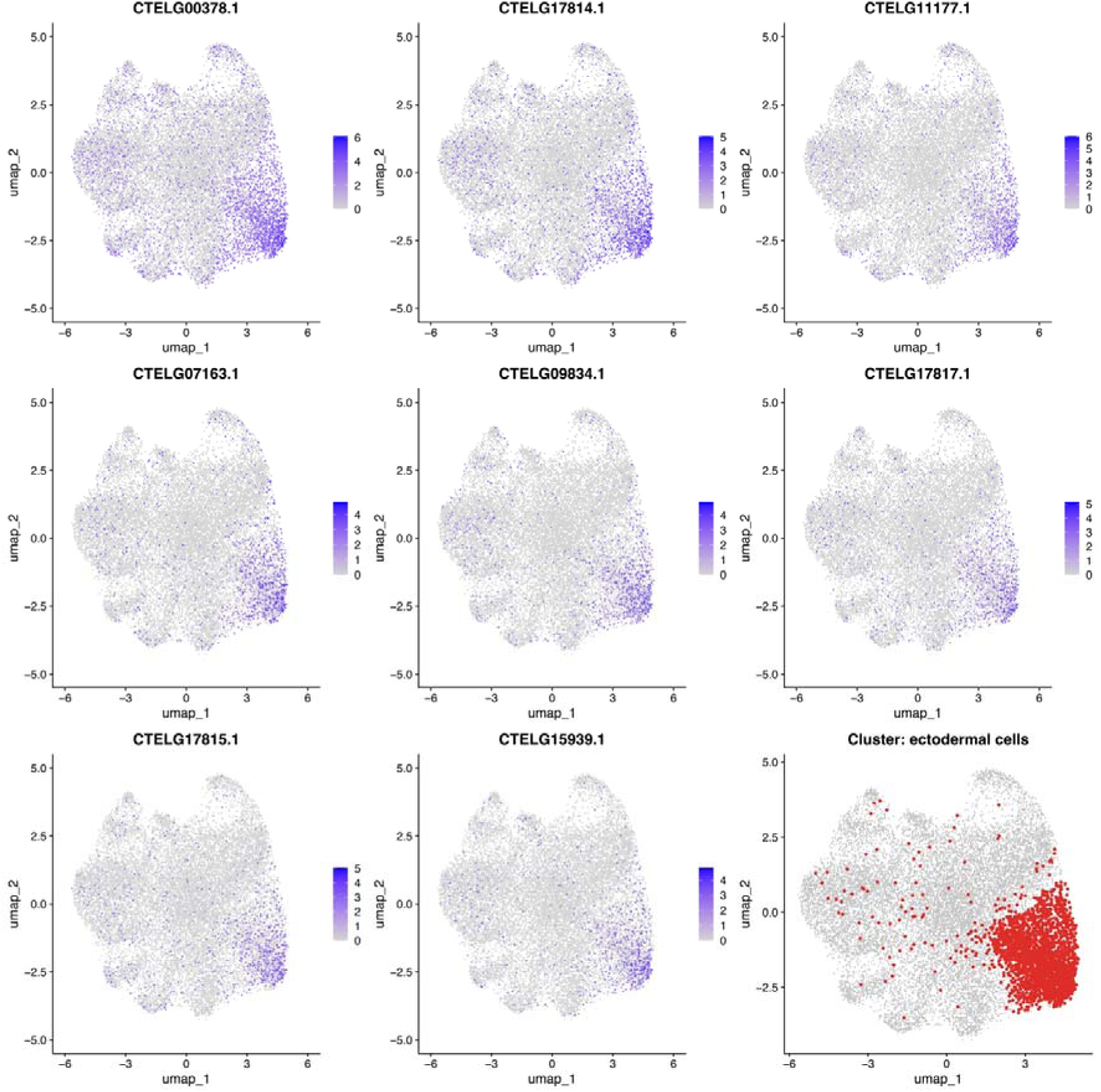
UMAP plots for the top eight marker genes for cluster 3 (ectodermal cells). UMAP plots are coloured based on expression levels obtained from the Wilcoxon Rank Sum differential expression test. The colour bar indicates the expression level (natural log) of each gene.

**Supplementary Figure 15.**
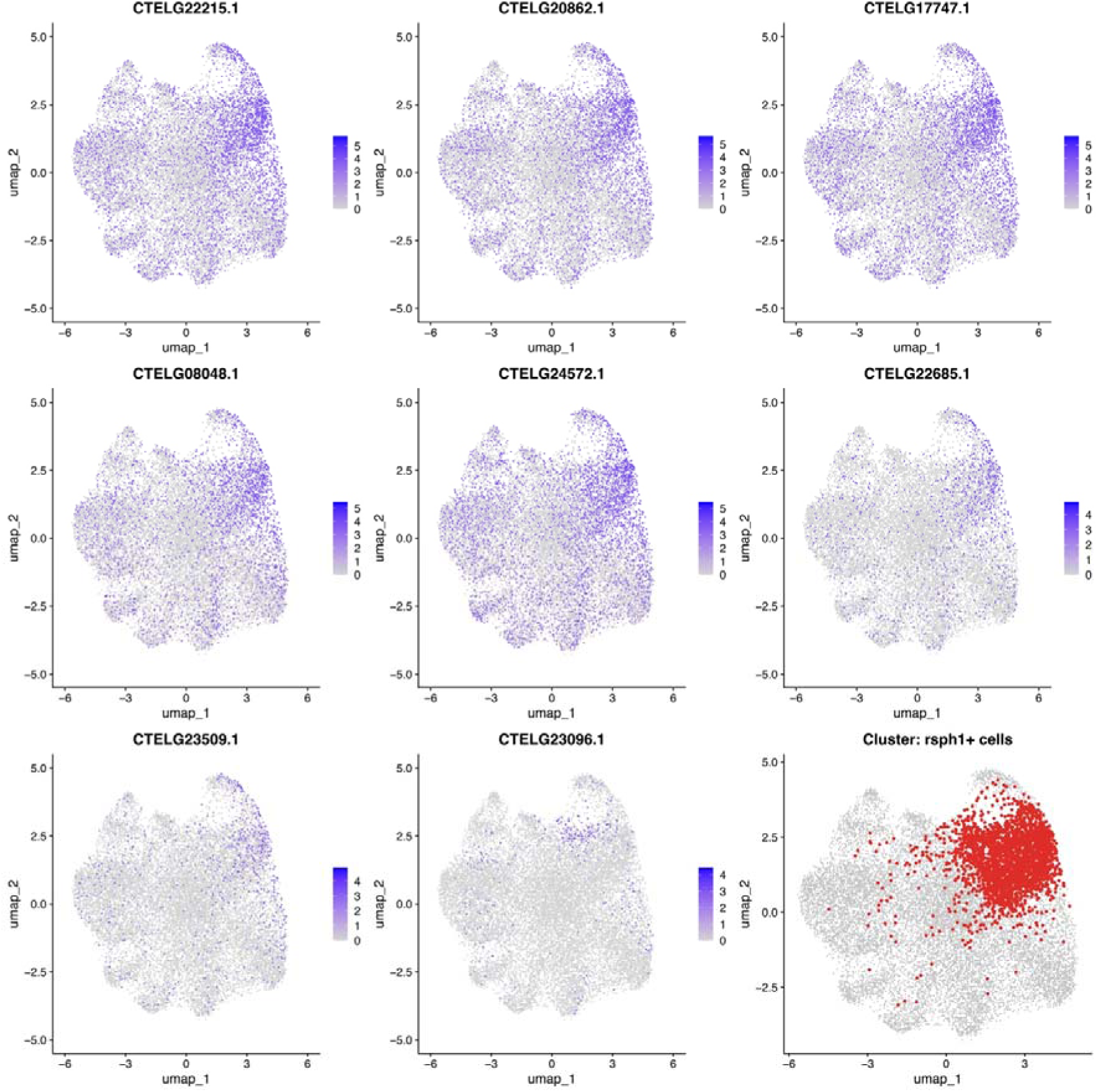
UMAP plots for the top eight marker genes for cluster 4 (*rsph1*^+^/tubulin^high^ cells). UMAP plots are coloured based on expression levels obtained from the Wilcoxon Rank Sum differential expression test. The colour bar indicates the expression level (natural log) of each gene.

**Supplementary Figure 16.**
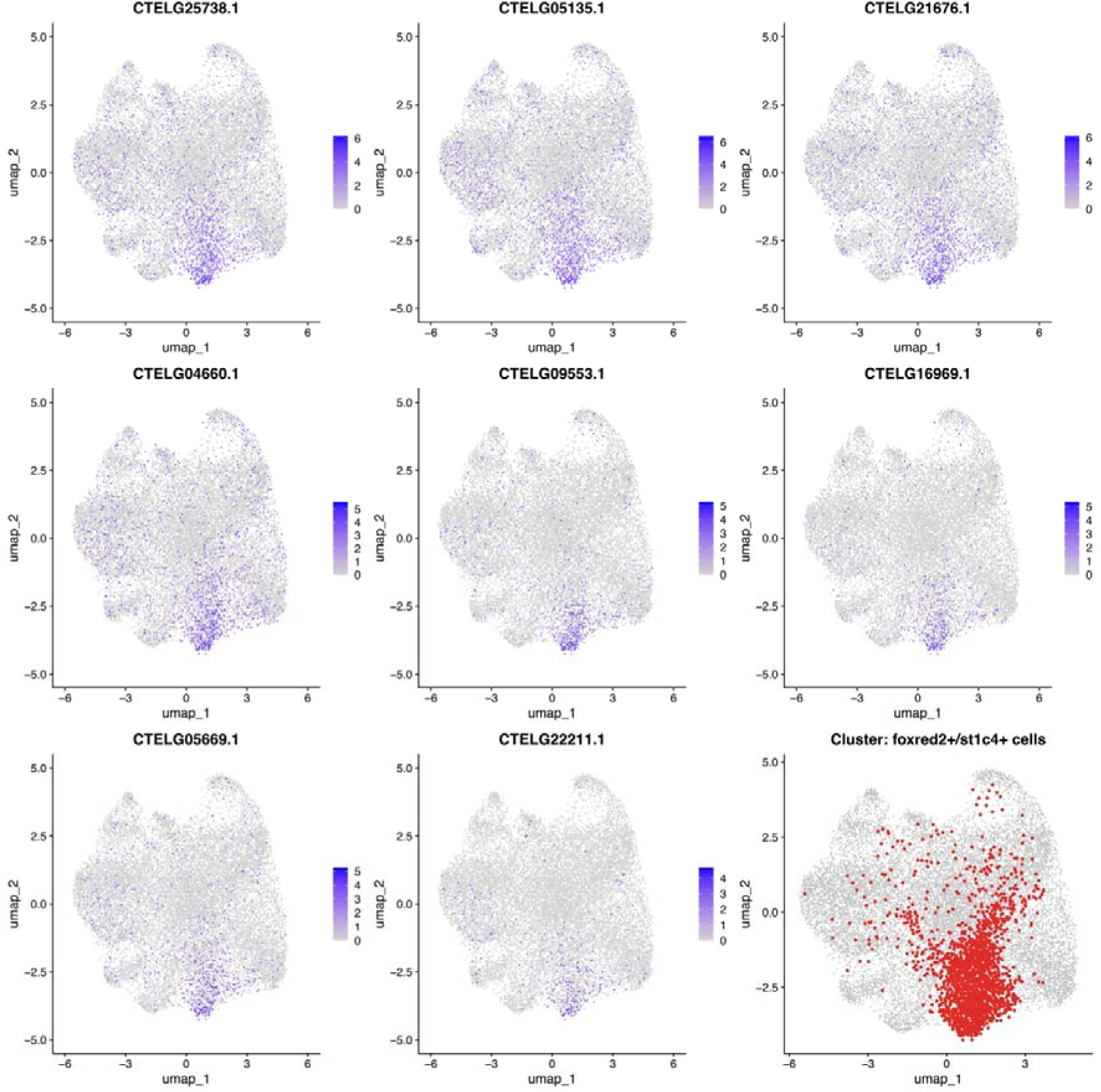
UMAP plots for the top eight marker genes for cluster 5 (*foxred2*^+^ cells). UMAP plots are coloured based on expression levels obtained from the Wilcoxon Rank Sum differential expression test. The colour bar indicates the expression level (natural log) of each gene.

**Supplementary Figure 17.**
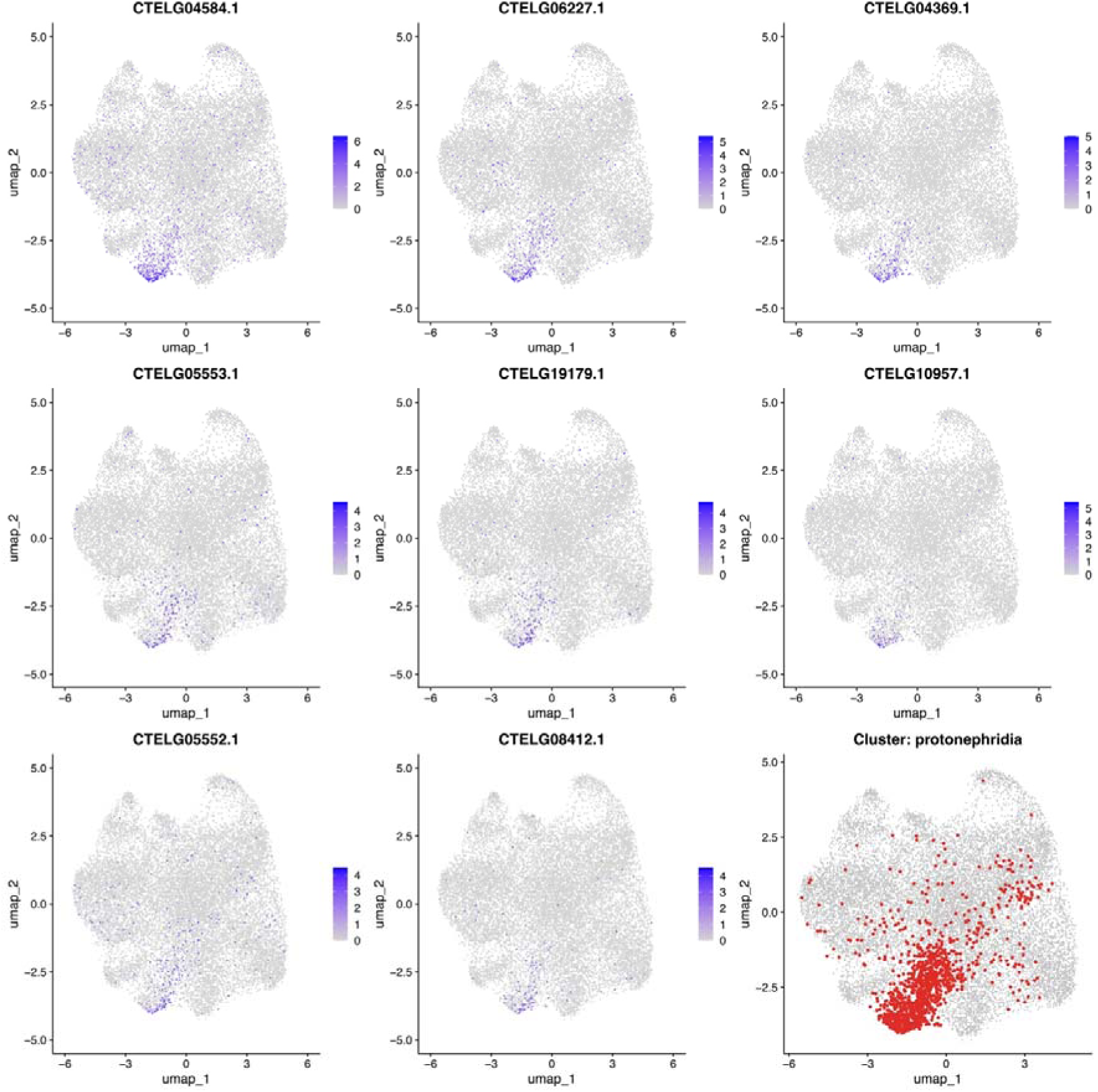
UMAP plots for the top eight marker genes for cluster 6 (protonephridia; ectodermal cells). UMAP plots are coloured based on expression levels obtained from the Wilcoxon Rank Sum differential expression test. The colour bar indicates the expression level (natural log) of each gene.

**Supplementary Figure 18.**
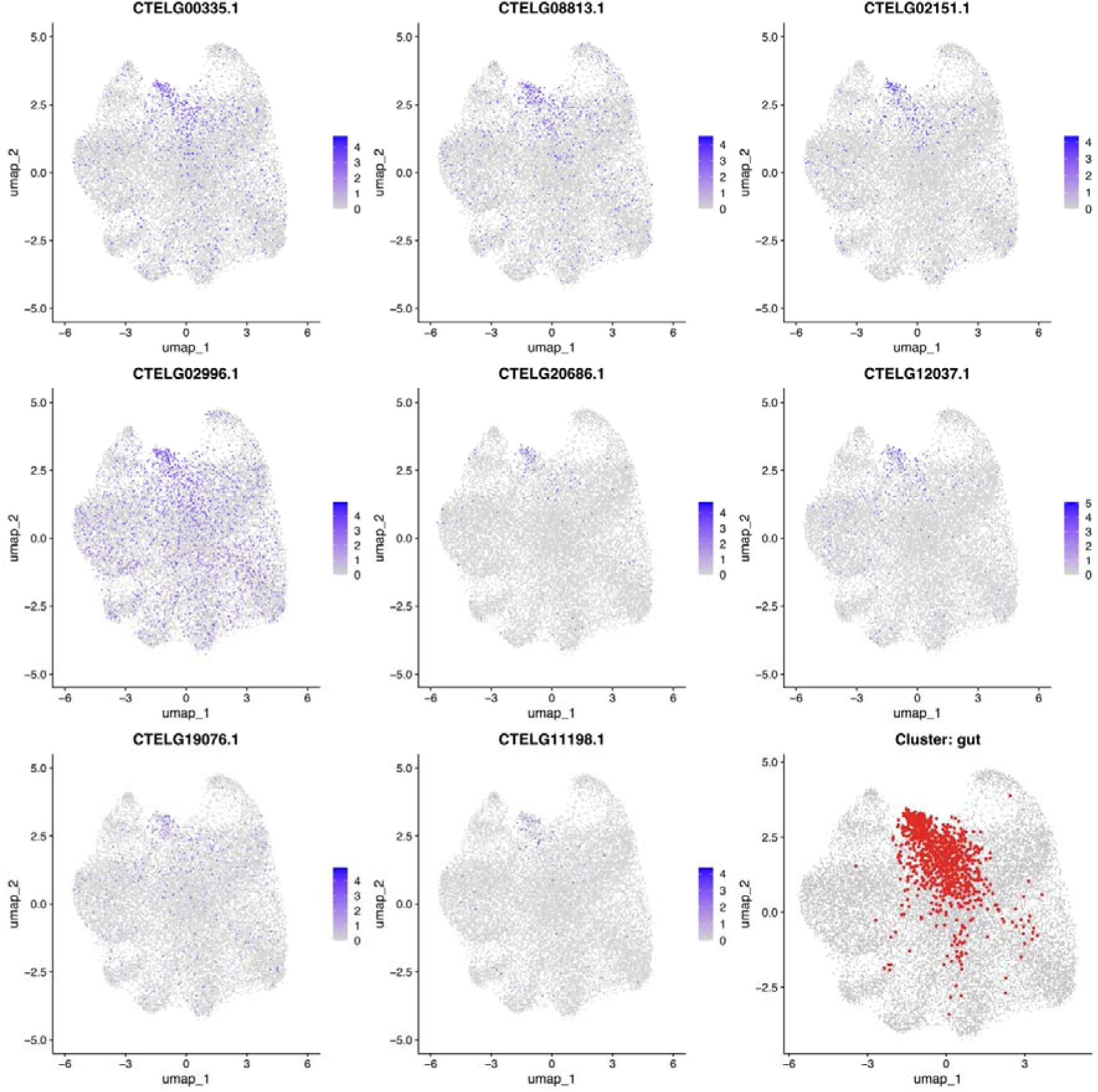
UMAP plots for the top eight marker genes for cluster 7 (gut). UMAP plots are coloured based on expression levels obtained from the Wilcoxon Rank Sum differential expression test. The colour bar indicates the expression level (natural log) of each gene.

**Supplementary Figure 19.**
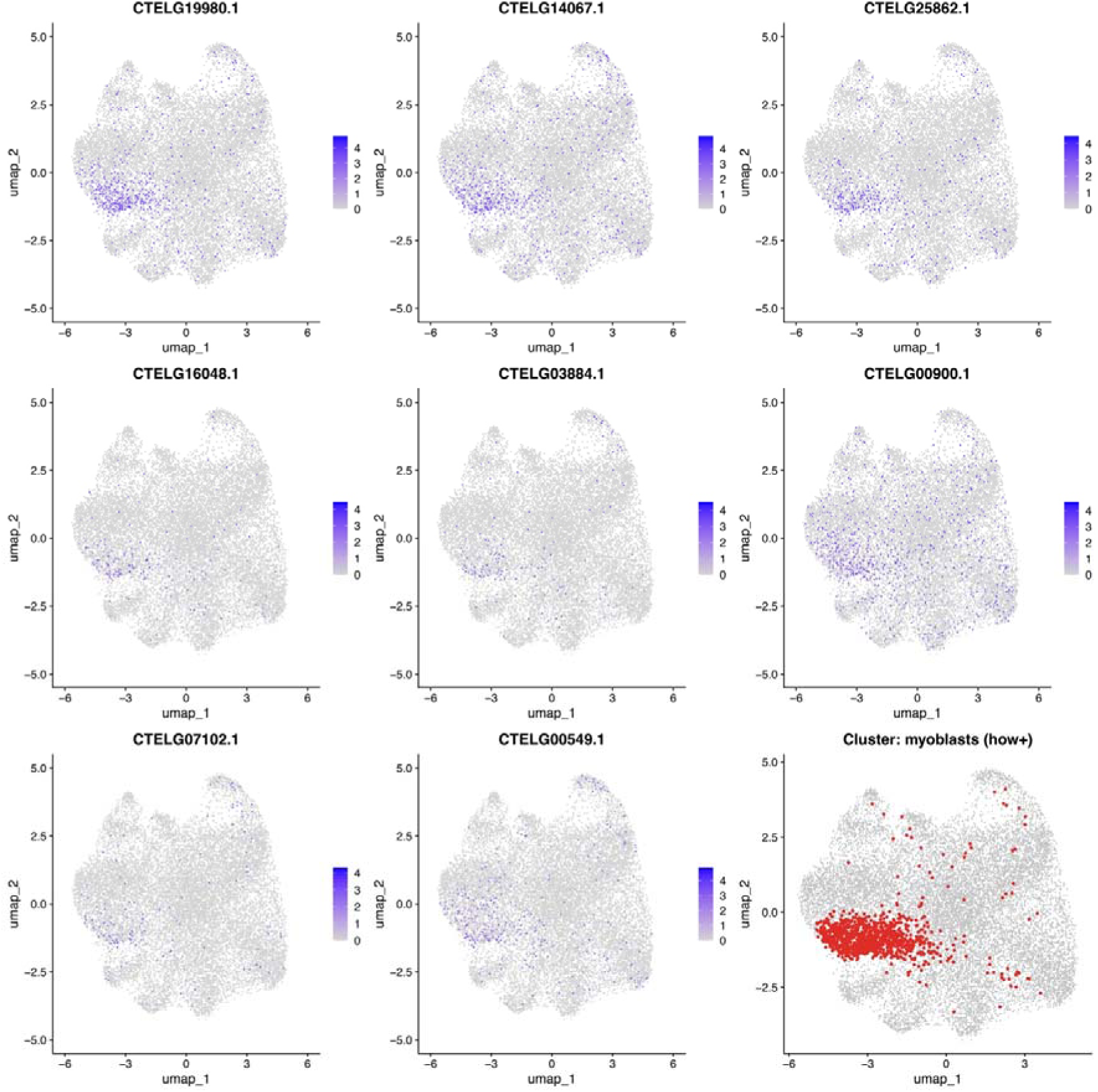
UMAP plots for the top eight marker genes for cluster 8 (myoblasts). UMAP plots are coloured based on expression levels obtained from the Wilcoxon Rank Sum differential expression test. The colour bar indicates the expression level (natural log) of each gene.

**Supplementary Figure 20.**
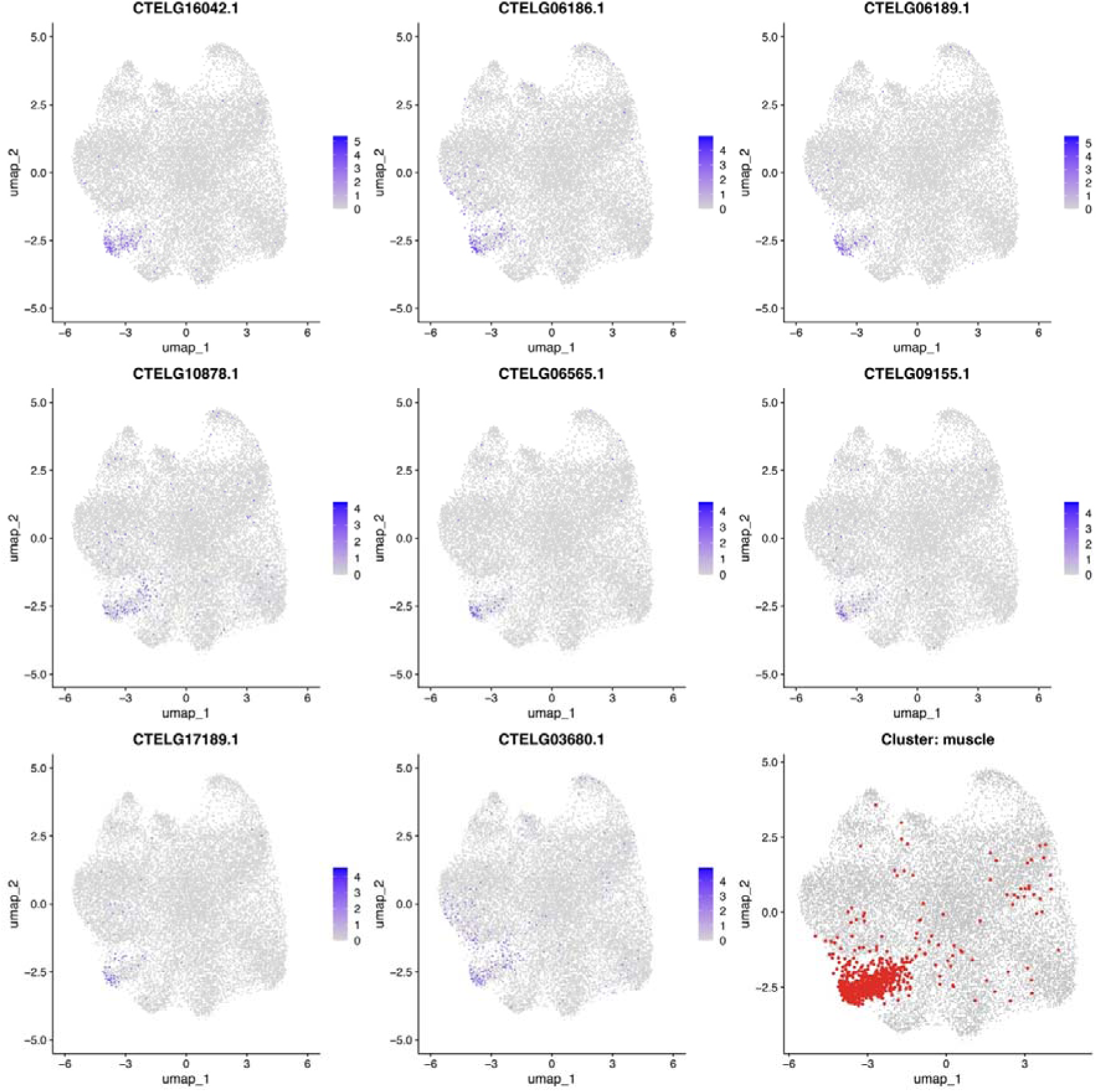
UMAP plots for the top eight marker genes for cluster 9 (muscle). UMAP plots are coloured based on expression levels obtained from the Wilcoxon Rank Sum differential expression test. The colour bar indicates the expression level (natural log) of each gene.

**Supplementary Figure 21.**
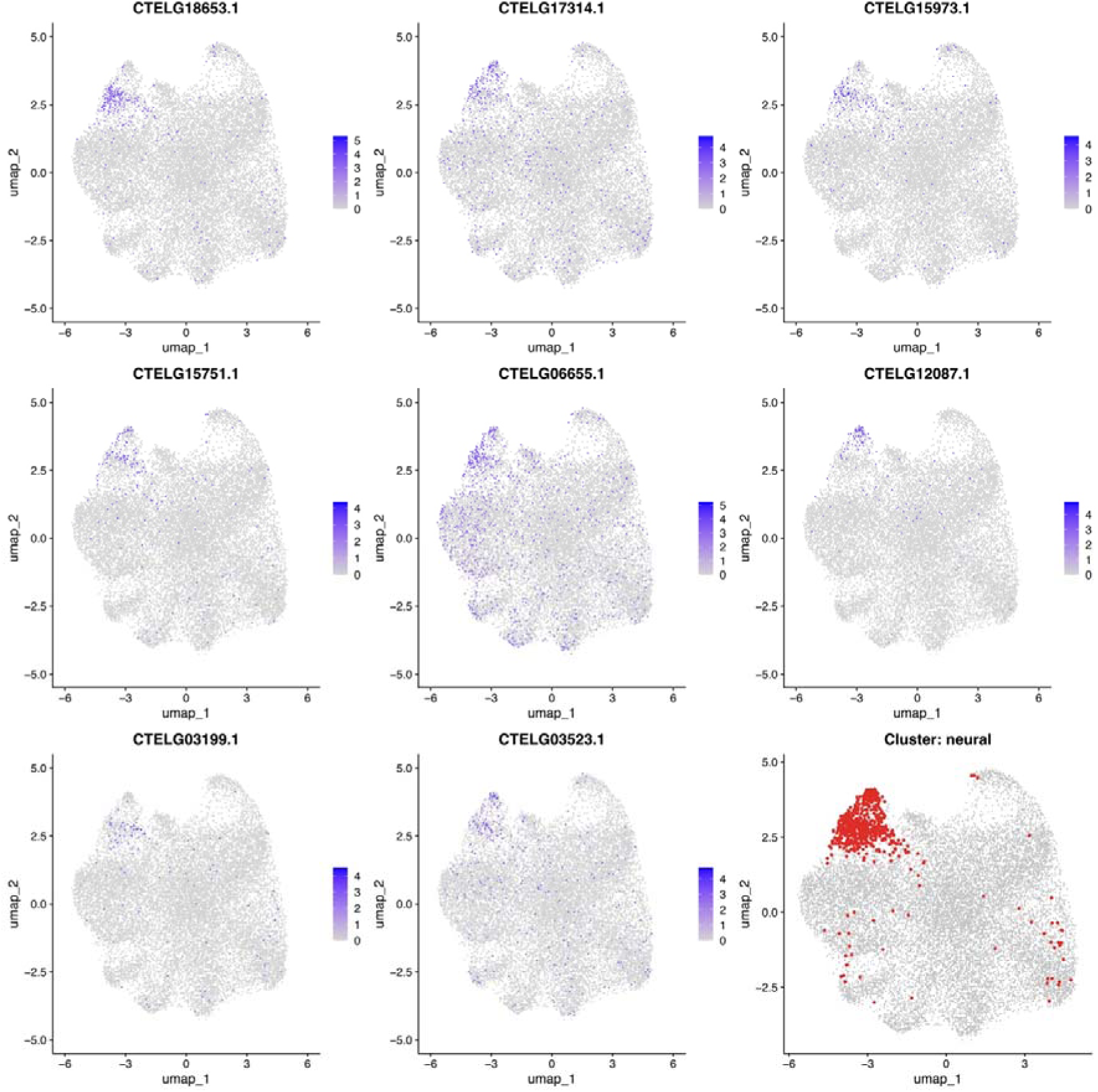
UMAP plots for the top eight marker genes for cluster 10 (neural). UMAP plots are coloured based on expression levels obtained from the Wilcoxon Rank Sum differential expression test. The colour bar indicates the expression level (natural log) of each gene.

**Supplementary Figure 22.**
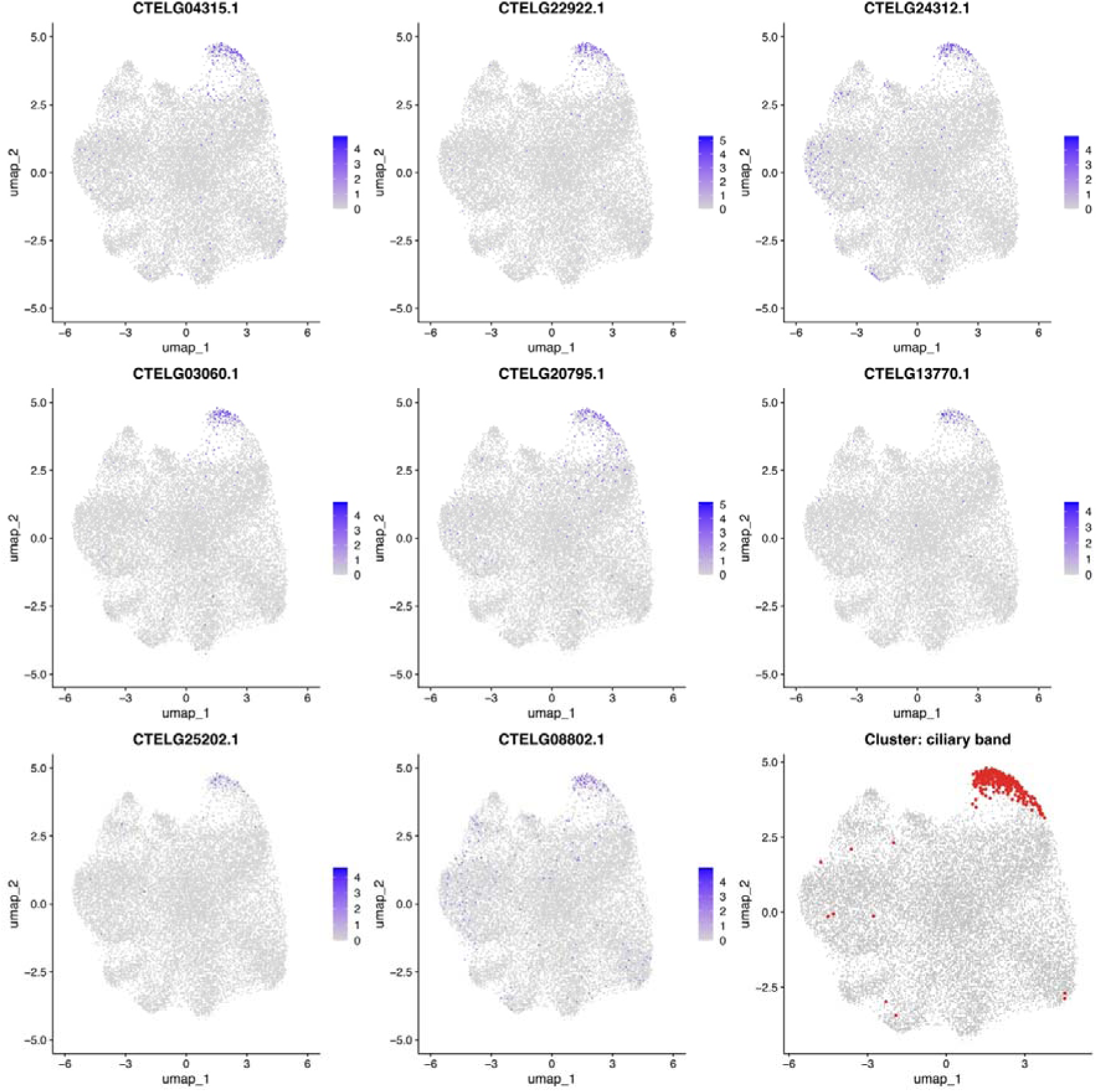
UMAP plots for the top eight marker genes for cluster 11 (ciliary band). UMAP plots are coloured based on expression levels obtained from the Wilcoxon Rank Sum differential expression test. The colour bar indicates the expression level (natural log) of each gene.

## Index of Supplementary Tables

**Supplementary Table 1.** Summary of sequenced long-read libraries from Magattract HMW gDNA extractions.

**Supplementary Table 2.** Summary of the repeat landscape in the genome of *C. teleta*.

**Supplementary Table 3.** List of genomes and annotations used in the gene family evolutionary analysis.

**Supplementary Table 4.** Orthogroup reconstruction.

**Supplementary Table 5.** Summary of the gene family evolutionary analysis.

**Supplementary Table 6.** List of orthogroups expanded in *C. teleta*.

## Index of Supplementary Files

**Supplementary File 1.** Alignment of *cox1* sequences used for phylogenetic reconstruction.

